# Distinct Rab11-associated membrane trafficking pathways bidirectionally control neuronal extracellular vesicle cargo traffic

**DOI:** 10.64898/2026.02.25.708063

**Authors:** Amy L. Scalera, Cassandra R. Blanchette, Erica C. Dresselhaus, Eric Gomez, Jack Y. Cheng, Avital A. Rodal

**Author notes:** These authors contributed equally (Amy L. Scalera, Cassandra R. Blanchette).

## Abstract

Neuronal extracellular vesicles (EVs) are released from synapses, and play roles in cellular communication, proteostasis, and the spread of toxic proteins in disease. The small GTPase Rab11 is required to maintain a reservoir of EV cargoes at presynaptic terminals, but how its diverse effector proteins contribute to this function and where Rab11 acts in neurons remains unclear. Using *Drosophila* motor neurons as a model, we show that EV cargoes redistribute from synapses to axons and cell bodies in *rab11* mutants, concomitant with reduced release from synapses. We conducted a directed genetic screen of Rab11-associated factors and found that they have distinct roles in EV trafficking. Tethering and sorting factors are required to maintain levels of presynaptic EV precursors, supporting the hypothesis that Rab11 regulates EV cargo pools through recycling flux rather than by directly mediating EV release. Unexpectedly, we found that different classes of Rab11-associated proteins have opposite functions: the motor protein MyoV and the PI4KIIIα component Rbo sustain cargo levels at synapses, while the motor adaptor Nuf/Rab11FIP4 and the PI4KIIIβ homolog Fwd restrict cargo levels. Together, these results indicate that Rab11 regulates multiple distinct organelle transport trajectories and PI(4)P populations to direct EV cargoes toward different cellular fates.

## Introduction

Neurons rely on a highly regulated system of endosomal compartments to sort and transport cargoes involved in signaling, growth, and disease. These dynamic and mobile membrane compartments mature and change identity, both over time and in response to cellular stimuli (Winckler et al., 2018; Yarwood et al., 2020). Compartment identity is regulated by Rab GTPases, which act as switches by cycling between an active GTP-bound state and an inactive GDP-bound state (Homma et al., 2021; Naslavsky & Caplan, 2018). In their active state, Rabs bind to effector proteins, regulating membrane remodeling and compartment mobility, altering membrane lipid and protein composition, and driving the maturation of endosomal compartments (Bouchet et al., 2018; Campa & Hirsch, 2017; Horgan & McCaffrey, 2009; Jing & Prekeris, 2009). Individual Rab family members are specifically targeted to distinct endosomal compartments, providing researchers with spatial and molecular experimental handles to interrogate the regulation of that compartment (Pfeffer, 2017).

In addition to their role in sorting and trafficking cargo within the cell, Rab GTPase-mediated regulation of endosomal compartments also facilitates the packaging of cargo into extracellular vesicles (EVs), which can be distinguished as microvesicles—released by direct budding from the plasma membrane—or exosomes, which are released when multivesicular bodies (MVBs) fuse with the plasma membrane (Blanchette & Rodal, 2020; van Niel et al., 2018). In neurons, EVs play roles in intercellular communication, proteostasis, and propagation of toxic proteins in neurodegenerative disease (Dresselhaus et al., 2024; Pitt et al., 2016; Quek & Hill, 2017). EV regulation is especially complex in neurons due to their unique polarized, elongated morphology and specialized membrane trafficking pathways. However, we have limited understanding of how transport and release of neuronal EV cargoes is controlled.

The *Drosophila* larval neuromuscular junction (NMJ) serves as an excellent *in vivo* model for studying synaptic EV trafficking. EVs released from presynaptic terminals are taken up by muscles or glia, or accumulate extracellularly within the folds of the postsynaptic muscle cell plasma membrane, allowing released cargoes to be readily visualized by confocal microscopy (Ashley et al., 2018; Blanchette et al., 2022; Fuentes-Medel et al., 2009; Koles et al., 2012; Korkut et al., 2009, 2013; Lauwers et al., 2018; Walsh et al., 2021). Multiple EV cargoes have been identified and characterized at the *Drosophila* NMJ, enabling parallel cell biological and functional studies (Blanchette et al., 2022; Koles et al., 2012; Korkut et al., 2013; Walsh et al., 2021). The majority of the observed EV cargoes in this system are likely packaged into exosomes rather than microvesicles, as synaptic MVBs harbor known EV cargoes, and the released EVs are the expected size for exosomes (Dresselhaus et al., 2024; Koles et al., 2012; Korkut et al., 2009; Walsh et al., 2021).

The Rab GTPase Rab11 is essential for directing cargo from early endosomes to the cell surface through recycling compartments (Akbergenova & Littleton, 2017; González-Gutiérrez et al., 2020; Sönnichsen et al., 2000), as well as for regulating exit of cargoes from the Golgi apparatus (Pantazopoulou et al., 2014; Pinar et al., 2015). Rab11 function is also crucial in regulating EV release, as disruptions to Rab11 lead to significant impairments in EV cargo transport, both in mammalian cell culture models and at the *Drosophila* NMJ (Ashley et al., 2018; Blanchette et al., 2022; Escudero et al., 2014; Koles et al., 2012; Korkut et al., 2013; Savina et al., 2002; Walsh et al., 2021). Specifically, in *Drosophila rab11* mutants, cargoes are depleted from NMJ presynaptic terminals, leading to a reduction in their release into the postsynaptic region via EVs and a loss of their physiological functions (Korkut et al., 2013; Walsh et al., 2021). Similarly, disrupting canonical endocytic proteins leads to a local depletion of synaptic EV cargoes, likely due to impaired trafficking between the plasma membrane and Rab11 recycling compartments and increased diversion into degradative pathways (Blanchette et al., 2022). Conversely, mutations in retromer, an endosomal sorting complex, lead to an accumulation of EV cargo proteins in Rab11-positive compartments and a concomitant increase in EV release (Walsh et al., 2021). These findings suggest that one or more steps of Rab11-dependent traffic are critical for EV cargo transport and function.

Several key questions remain regarding how Rab11-dependent cargo sorting influences EV transport. First, it is unclear whether Rab11 governs EV biogenesis through its canonical roles in endosome-to-plasma membrane recycling or Golgi exit (e.g. by indirectly regulating loading of cargoes into EV precursors), or through a unique function in EV trafficking (e.g. by directly promoting fusion of MVBs with the plasma membrane (Koles et al., 2012; Messenger et al., 2018; Savina et al., 2005)). Second, it is not yet determined whether Rab11-mediated EV cargo trafficking events occur locally at synapses or involve long-distance transport between the cell body and synapses. Finally, the specific Rab11 effectors involved in EV biogenesis and the nature of trafficking events that they regulate remain unknown. These effectors may shape the stepwise progression of EV biogenesis via distinct pathways, including facilitating Golgi exit, MVB formation, activating motor proteins for endosome transport, or tethering endosomes to the plasma membrane. Here, we investigate how Rab11 regulates neuronal EV cargo trafficking by dissecting the contributions of its binding partners, effectors, and other regulators of endosomal recycling, and determining where these components act within the complex, polarized architecture of neurons.

## Results

### Rab11 regulates EV cargo distribution between synapses, axons, and cell bodies

To investigate the previously reported reduction of EV cargoes in presynaptic terminals of *rab11* loss-of-function mutants (Koles et al., 2012; Korkut et al., 2013; Walsh et al., 2021), we examined whether EV cargo levels were also altered in other regions of the neuron (**Fig. 1A**). To this end, we quantified levels of the EV cargoes Amyloid Precursor Protein-GFP (APP, GAL4-UAS driven) (**Fig. 1B-F**) and Synaptotagmin-4-GFP (Syt4, endogenously tagged) (**Fig. 1G-I**) in cell bodies, the central neuropil (which contains motor neuron dendrites), axons, and the NMJ.

**Figure 1.**
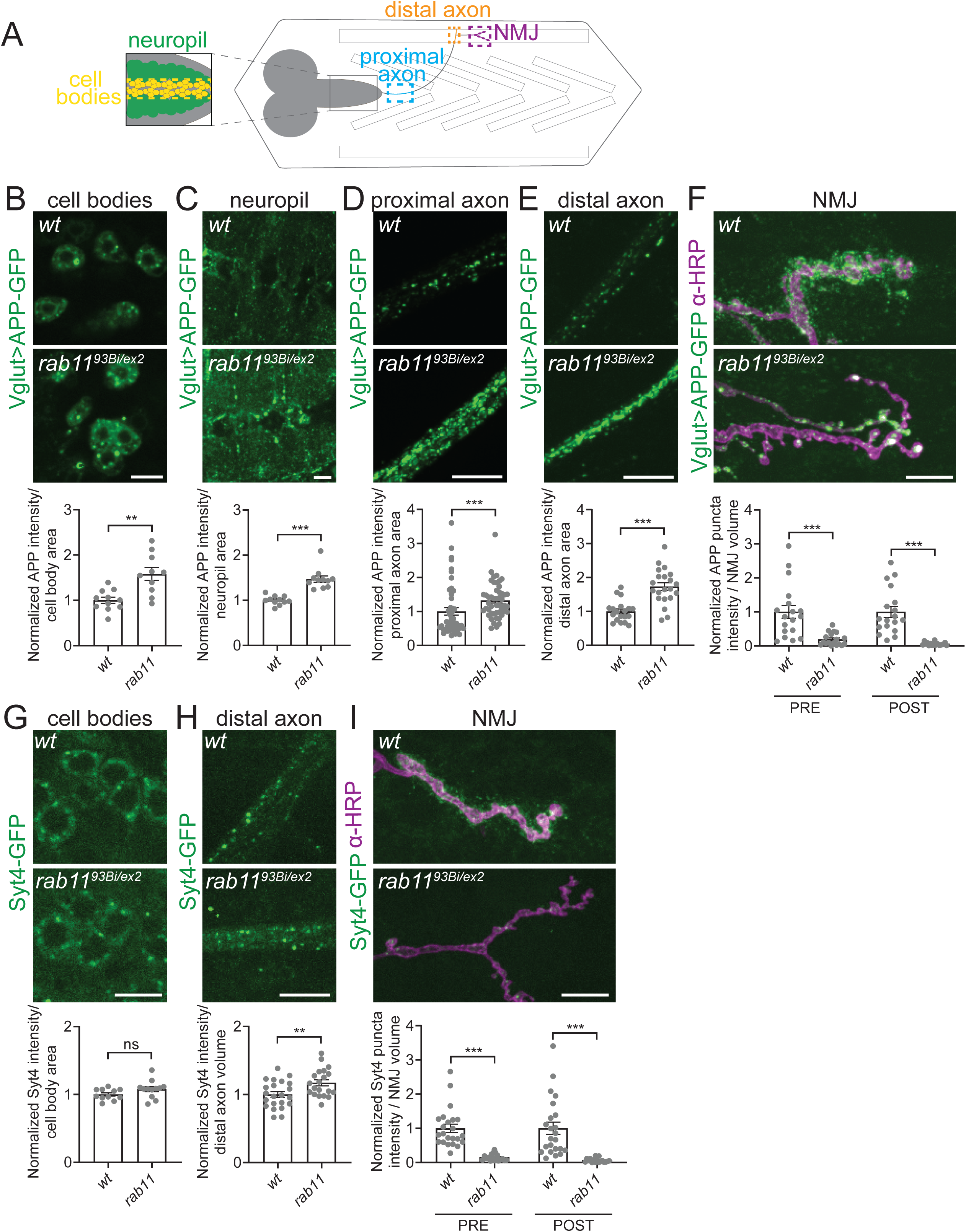
Rab11 regulates EV cargo distribution between synapses, axons, and cell bodies. **(A)** Diagram of *Drosophila* third instar larva highlights regions of the nervous system in which EV cargoes were imaged and quantified: motor neuron cell bodies in the ventral nerve cord (yellow), neuropil (green), proximal axons projecting from motor neuron cell bodies (blue), distal axons adjacent to the NMJ (orange), and neuromuscular junctions (purple). **(B-F)** Loss of *rab11* leads to an increase in APP-GFP levels in cell bodies, the neuropil, and proximal/distal axons, along with depletion from NMJs. **(B)** Representative images of APP-GFP in cell bodies and quantification of mean intensity. **(C)** Representative images of APP-GFP in the neuropil and quantification of mean intensity. **(D)** Representative images of APP-GFP in proximal axons and quantification of mean intensity. **(E)** Representative images of APP-GFP in distal axons and quantification of mean intensity. **(F)** Representative images of APP-GFP at muscle 6/7 NMJs and quantification of pre- and postsynaptic APP-GFP puncta intensity. **(G-I)** Loss of *rab11* leads to an increase in Syt4-GFP in axons along with depletion from NMJs. **(G)** Representative images of Syt4-GFP in cell bodies and quantification of mean intensity. **(H)** Representative images of Syt4-GFP in distal axons and quantification of mean intensity. **(I)** Representative images of Syt4-GFP at muscle 4 NMJs and quantification of pre- and postsynaptic Syt4-GFP puncta intensity. Data is represented as mean ± SEM; n is depicted by individual gray dots in the graphs and represents brains **(B,C,G)**, axons **(D,E,H)** or NMJs **(F,I)**. Cell body, neuropil, and axon intensity measurements were normalized to area or volume of compartment (as noted in graphs), NMJ intensity measurements were normalized to presynaptic volume; all measurements were further normalized to the mean of their respective controls. All scale bars are 10 µm. See **Tables S1** and **S2** for detailed genotypes, sample sizes, and statistical analyses. wt, wild-type. α-HRP was used to label neuronal membranes.

We first measured APP-GFP and Syt4-GFP levels in the motor neuron ventral ganglion. In *rab11* hypomorphs, APP-GFP mean intensity was significantly increased in both cell bodies (**Fig. 1B**) and the neuropil (**Fig. 1C**) compared to controls, while Syt4-GFP levels in cell bodies remained unchanged (**Fig. 1G**). Next, in the axonal region within 50 µm of the ventral ganglion (proximal axon), we found that APP-GFP levels were significantly increased in *rab11* hypomorphs compared to controls (**Fig. 1D)**. We then measured APP-GFP and Syt4-GFP levels in the axonal region immediately adjacent to the NMJ (distal axon), and found that both were significantly elevated (**Fig 1E, H**). Finally, as previously observed, we found a dramatic reduction of both EV cargoes in the presynaptic region of the NMJ, with a concomitant reduction in their levels in EVs that had been released into the postsynaptic region (**Fig. 1F,I**).

To assess whether these changes reflected overall protein level changes or simply cargo redistribution, we estimated the total amount of Syt4-GFP within the neuron. Using our measured mean intensities from these regions and estimates of compartment volume (see Materials and Methods), we found that the total amount of Syt4-GFP in the neuron was similar in *rab11* hypomorphs (85.7 +/−8.0 normalized fluorescence units/µm^3^) compared to controls (100 +/- 16.2 normalized fluorescence units/µm^3^) (n=5; Student’s t-test p=0.11). In *rab11* hypomorphs, our estimates indicate that 44% of the Syt4-GFP was found in in the cell body, 43% in axons, and 13% at the NMJ, compared to 34% in cell bodies, 33% in axons, and 33% at the NMJ in control animals. These findings suggest that loss of *rab11* causes a spatial redistribution of EV cargoes from synaptic terminals to axons and cell bodies.

### Stalled EV cargo-containing vesicles are increased in *rab11* mutant axons

To determine whether axonal accumulation of EV cargoes resulted from altered transport dynamics, we performed live imaging of motor neuron-driven APP-GFP in axons proximal to the cell body. Kymograph analysis revealed no differences in the number of anterograde, retrograde, or complex tracks between *rab11* mutants and controls (**Fig. 2A,B**). Additionally, no significant changes were detected in the velocity of APP-GFP puncta in either direction (**Fig. 2C**), indicating that overall cargo movement dynamics were unaffected. However, we observed a significant increase in the number of stationary APP-GFP puncta in *rab11* mutants (**Fig. 2A,B**). Collectively, these findings suggest that the increased axonal EV cargoes observed in *rab11* mutants result from an accumulation of stationary compartments rather than enhanced trafficking in the anterograde or retrograde directions. Further, since we found no defects in transport rates or directionality that would explain cargo depletion at the NMJ, our results suggest that the primary defect originates at the synapse itself. Therefore, we next focused on investigating Rab11’s synaptic activities to elucidate the specific mechanisms driving EV cargo depletion at these terminals.

**Figure 2.**
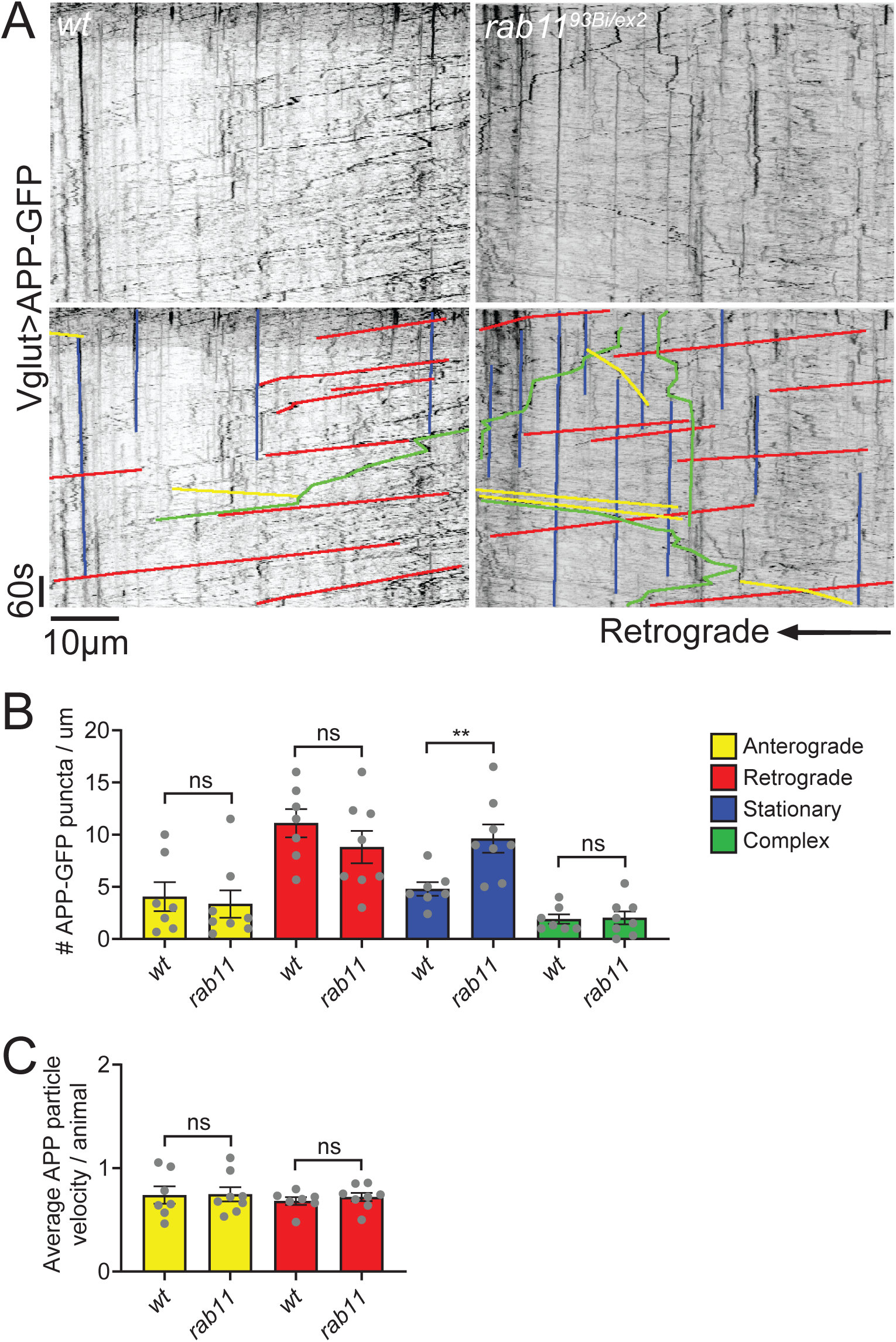
Stationary EV cargo-containing compartments are increased in *rab11* mutant axons. **(A)** (Top) Representative kymographs of live APP-GFP dynamics in control and *rab11* mutants. (Bottom) Overlay with colored lines indicating the direction of movement: anterograde (yellow), retrograde (red), stationary (blue), or complex (green). Brightness and contrast were adjusted separately for each image. **(B)** Quantification of average number of APP-GFP puncta moving in each direction per animal in control and *rab11* axons, demonstrating an increase in stationary APP-GFP puncta in *rab11* mutants. **(C)** Quantification of the average velocity of APP-GFP puncta undergoing anterograde or retrograde transport in control and *rab11* mutant axons. Data is represented as mean ± SEM; n is depicted by individual gray dots in the graphs and represents average value per animal (see **Methods** for details). See **Tables S1** and **S2** for detailed genotypes, sample sizes, and statistical analyses. wt, wild-type.

### A directed screen of Rab11 effectors, interactors, and other recycling machinery for roles in EV traffic

To unravel the mechanisms by which Rab11 maintains EV cargo levels at synapses, we took a candidate-based approach to investigate the roles of known and putative Rab11 effectors and binding partners, as well as regulators of endosomal recycling in EV traffic. To identify potential new effectors, we leveraged previously published proteomics data from *Drosophila* S2 cells, which identified interactors that bind specifically to Rab11 (Gillingham et al., 2014), and then focused on those with established synaptic functions. From this list of candidate genes, we collected and validated loss-of-function mutants or RNAi lines (**Table 1**), and tested their role in EV trafficking by measuring pre- and post-synaptic levels of the EV cargoes Neuroglian (Nrg, using a monoclonal antibody) or Syt4 (using our endogenous GFP knockin) (Walsh et al., 2021).

**Table 1.**
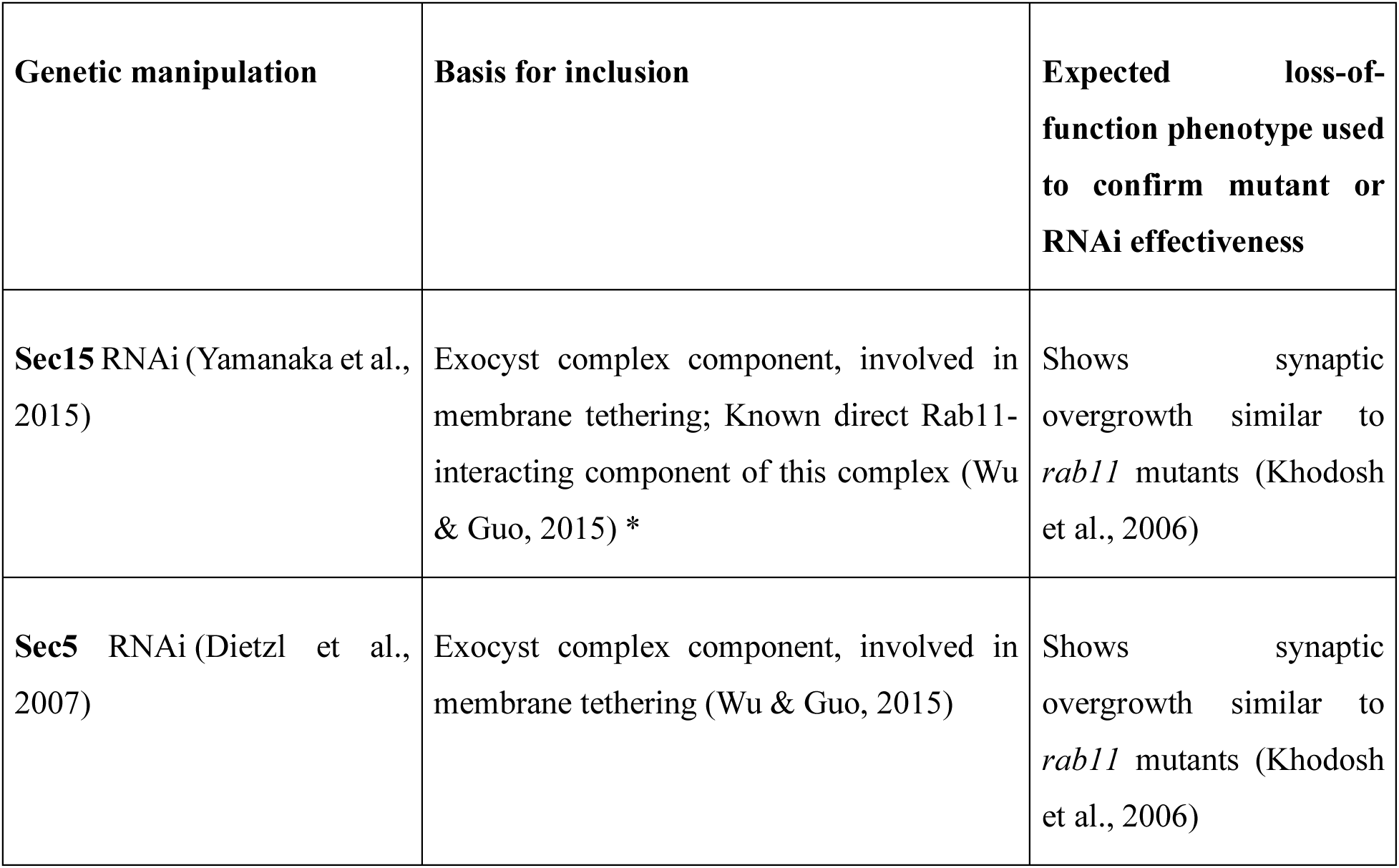
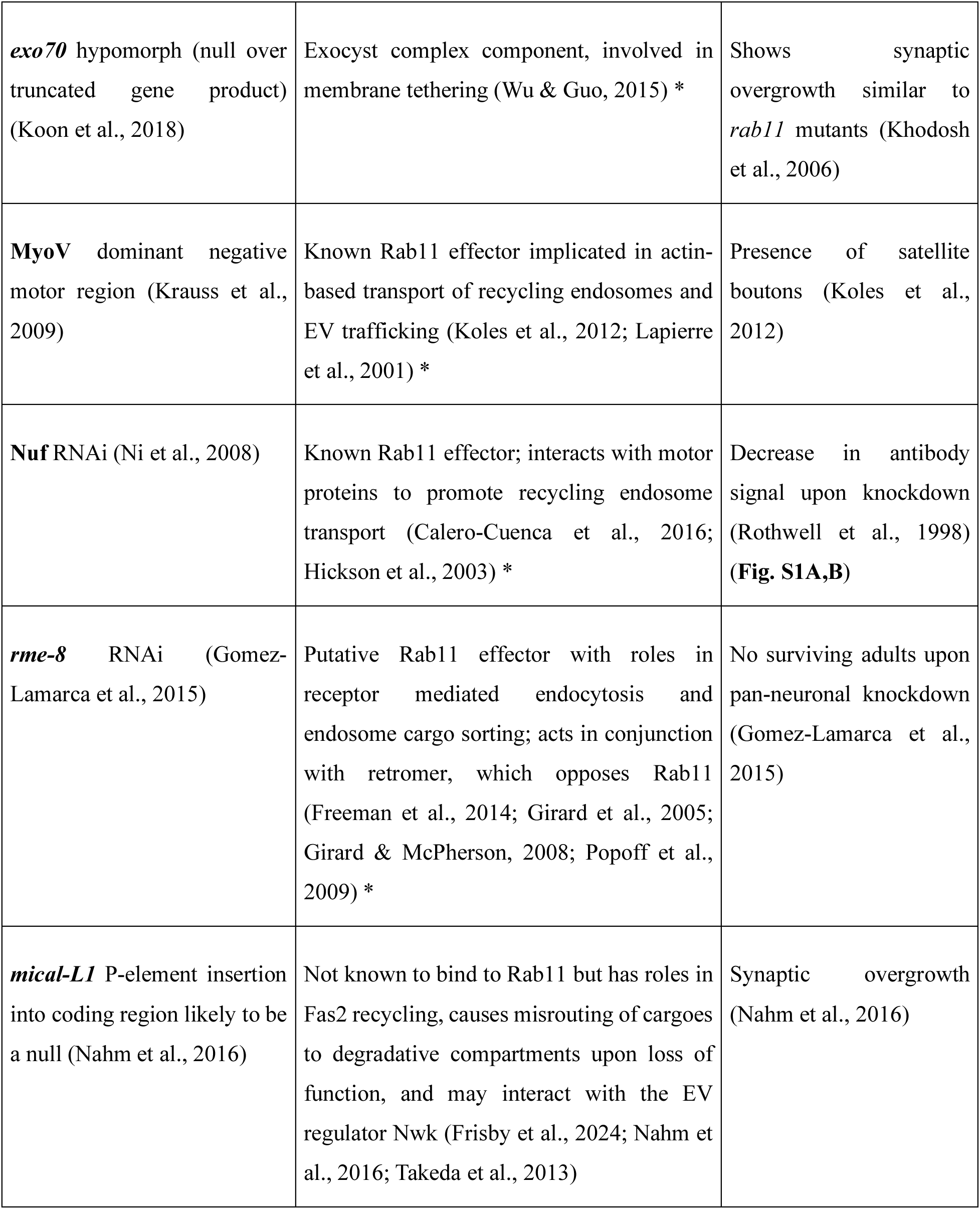
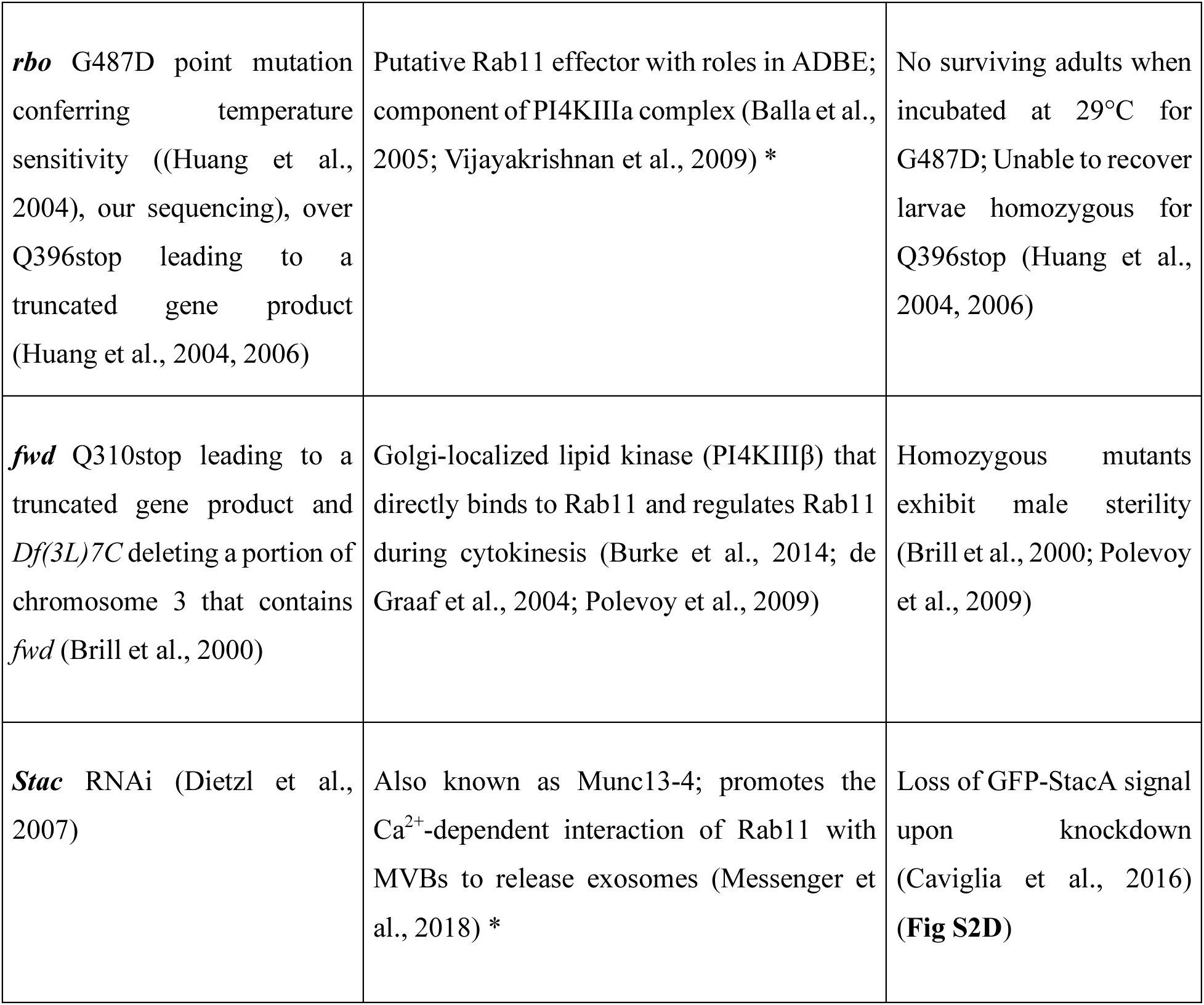
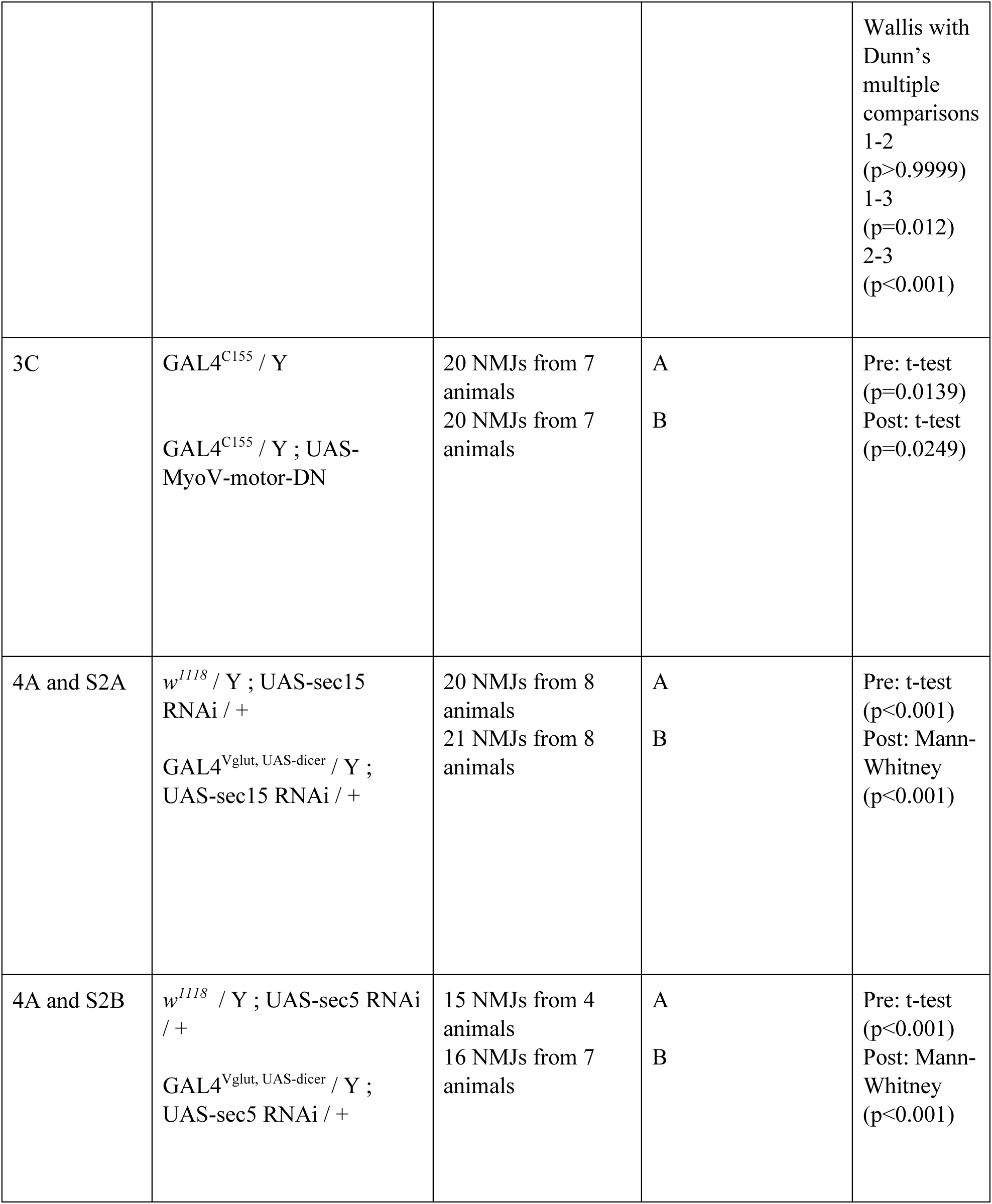
Recycling machinery candidates tested for roles in EV trafficking. List of genes tested for roles in EV trafficking, with justification for selection and phenotypic criteria used to validate each mutant or RNAi intervention. **\*** indicates Rab11-interactor identified in (Gillingham et al., 2014).

### Opposite functions of Rab11-interacting motor proteins and motor-regulatory factors in EV trafficking

Rab11 interacts with various motor proteins and their regulators to control membrane trafficking, but the roles of these specific effectors in EV biogenesis and transport are unknown. We hypothesized that motor-associated effectors of Rab11 might facilitate EV trafficking through two potential mechanisms: by promoting cargo transport to or from recycling endosomes to enhance EV cargo flux between the plasma membrane and endosomes, and/or by directly influencing MVB transport to the plasma membrane for fusion and intraluminal vesicle release. (**Fig. 3A**). To distinguish between these possibilities, we examined two key classes of Rab11 motor-associated effectors: Rab11 Family-Interacting Proteins (FIPs) and Myosin V (MyoV). If these effectors primarily function in mediating MVB-plasma membrane fusion, their disruption would result in decreased postsynaptic EV cargo levels accompanied by presynaptic cargo accumulation, as fusion-incompetent MVBs would be retained within presynaptic terminals (as has previously been observed for Hsp90 mutants (Lauwers et al., 2018)). However, if these effectors are primarily involved in the flux of EV cargoes through the recycling pathway, we would expect mutations that reduce flux to decrease EV cargo levels at both presynaptic and postsynaptic sites, while mutations that enhance flux would increase cargo levels at both sites (**Fig. 3A**).

**Figure 3.**
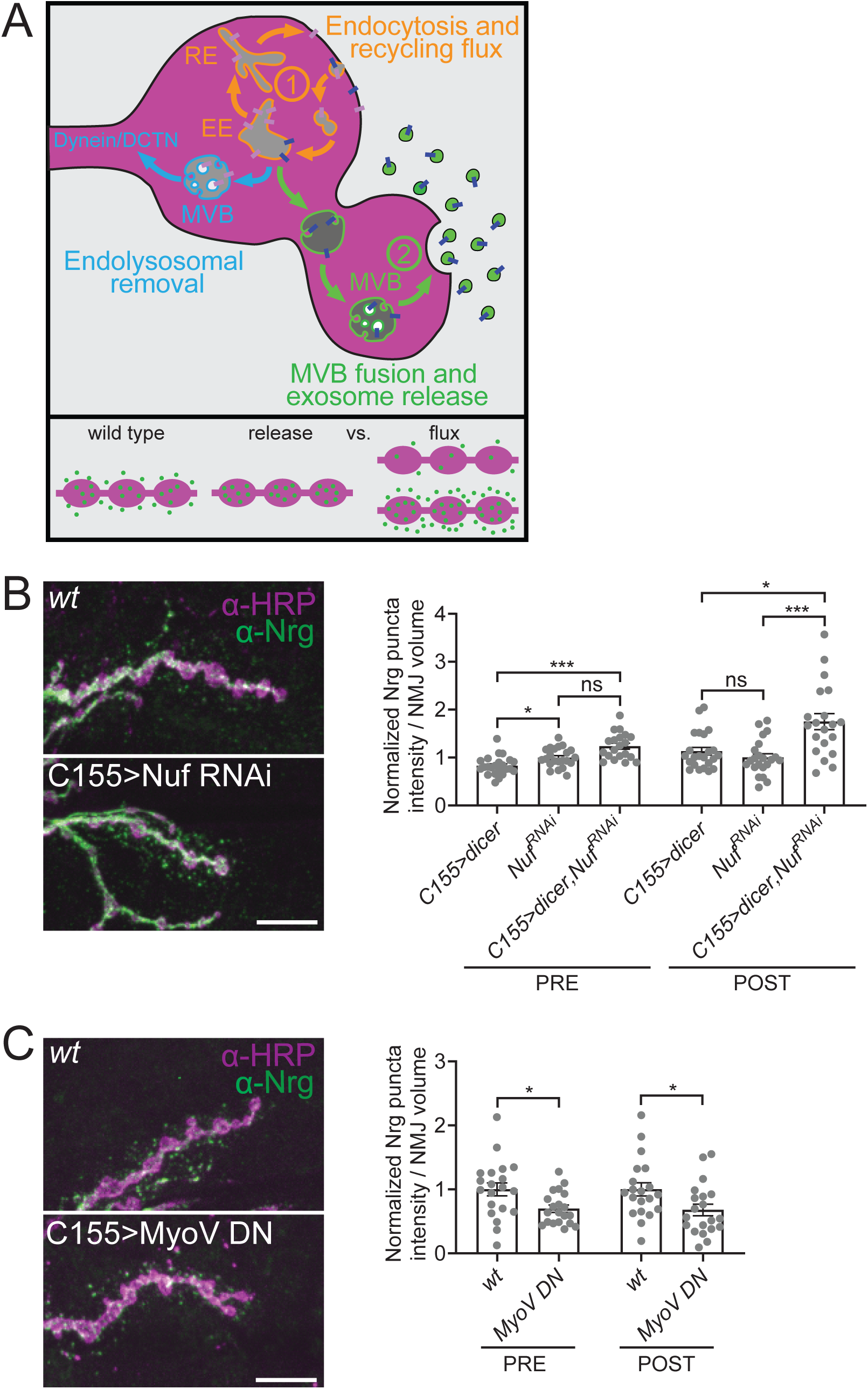
Opposite roles of Rab11-interacting motor proteins and motor regulatory factors in EV trafficking. **(A**) Model depicting alternative functions for Rab11 in recycling flux or MVB fusion, and predicted phenotypes for each mechanism. **(B)** Loss of *nuf* leads to an increase in Nrg at NMJs. Representative images of Nrg at muscle 6/7 NMJs and quantification of pre- and postsynaptic Nrg puncta intensity. **(C)** Disruption of MyoV leads to depletion of Nrg at NMJs. Representative images of Nrg at muscle 6/7 NMJs and quantification of pre- and postsynaptic Nrg puncta intensity. Data is represented as mean ± SEM; n is depicted by individual gray dots in the graphs and represents NMJs. NMJ intensity measurements were normalized to presynaptic volume; all measurements were further normalized to the mean of their respective controls. All scale bars are 10 µm. See **Tables S1** and **S2** for detailed genotypes, sample sizes, and statistical analyses. wt, wild-type. α-HRP was used to label neuronal membranes.

There are two classes of Rab11-FIPs, both of which have a 20 amino acid Rab11 binding site on their C-terminus but vary in their N-termini, with class I FIPs containing a phospholipid-binding domain and class II FIPs containing proline-rich and Ca^2+^-binding domains (Hales et al., 2001). *Drosophila* have a single Class I FIP, *dRip11*, which mediates entry of cargo into recycling endosomes (O’Sullivan & Lindsay, 2020; Prekeris et al., 2000; Schonteich et al., 2008). However, we were unable to detect specific α-dRip11 antibody signal in motor neurons to validate available RNAi tools, leaving the role of *dRip11* in neuronal EV trafficking undetermined. Next, we examined the single *Drosophila* Class II FIP, *nuclear fallout* (*nuf*), which is homologous to mammalian Rab11 FIP3 and FIP4 (Calero-Cuenca et al., 2016; Riggs et al., 2007). *nuf* has been implicated in motor-based transport and physically interacts with both Dynein and Rab11 (Riggs et al., 2003, 2007). We confirmed that Nuf is expressed in neurons and validated *nuf* knockdown by RNAi using α-Nuf antibody staining (**Fig. S1A**). Upon knockdown of *nuf*, we observed a striking increase in Nrg levels at both pre- and postsynaptic sites compared to controls (**Fig. 3B**), suggesting that Nuf normally acts to restrain EV release. This result is notable because it contrasts with the *rab11* mutant phenotype, indicating that Rab11 may, through its effectors, regulate independent pathways that either promote or limit the trafficking of cargoes into EV-precursor compartments.

We next tested the established Rab11 effector, Myosin V (MyoV), which interacts with Rab11 via its C-terminal tail and mediates short-range transport of recycling endosomes along actin filaments (Lapierre et al., 2001; Roland et al., 2011). Previous studies at the *Drosophila* NMJ demonstrated that MyoV disruption causes a reduction in the postsynaptic-to-presynaptic ratio of the EV cargo *Evenness interrupted* (Evi), leading to the hypothesis that Rab11 directs MyoV to transport MVBs for fusion with the plasma membrane (Koles et al., 2012). We investigated absolute changes in pre- and postsynaptic EV cargo levels upon neuronal expression of a dominant-negative form of MyoV (MyoV^DN^), which disrupts the motor activity of the protein (Krauss et al., 2009). In MyoV^DN^-expressing NMJs, we observed a significant decrease in Nrg levels at both pre- and postsynaptic sites compared to controls (**Fig. 3C**). These findings argue against the hypothesis that Rab11 primarily functions in MVB-plasma membrane fusion at the NMJ. Instead, they support a model in which Rab11-mediated recycling flux plays an important role in regulating EV cargo trafficking. In this model, MyoV promotes local EV cargo flux at synapses while Nuf restricts this traffic or facilitates cargo removal, suggesting bidirectional control of synaptic EV distribution by Rab11.

### Tethering effector proteins indicate a primary role for Rab11 in recycling flux of EV cargoes

To further test whether Rab11 functions primarily in recycling flux versus MVB fusion, we examined the role of tethering effectors in EV cargo traffic, beginning with the Exocyst complex. Exocyst is an eight-protein assembly that functions as a Rab11 effector, and is primarily known for tethering secretory vesicles to the plasma membrane to enable SNARE-dependent fusion (He & Guo, 2009; Lee et al., 2025). Similar to MyoV, Exocyst has been proposed to directly mediate Rab11-dependent MVB-plasma membrane fusion (Bai et al., 2022), Notably, previous work at the *Drosophila* NMJ showed that knockdown of *sec15* led to reduced postsynaptic levels of both Nrg and glycosylated EV cargoes marked by anti-HRP antibodies, but did not assess whether cargoes were trapped presynaptically as would be predicted (Kang et al., 2024). We tested this hypotheses by manipulating three Exocyst components: Sec5 (a core complex component), Sec15 (which binds Rab11), and Exo70 (which interacts with the membrane) (Wu & Guo, 2015). RNAi-mediated knockdown of Sec5 or Sec15, as well as a hypomorphic *exo70* mutation, led to decreased Nrg levels both pre- and postsynaptically (**Fig. 4A**; **Fig. S2A-C**). These findings, along with our MyoV results, argue against a primary role for Rab11 and its effectors in MVB-plasma membrane fusion. Instead they suggest that Rab11 and Exocyst collaborate to facilitate EV trafficking by promoting recycling compartment flux, ensuring a sufficient pool of cargoes for MVB loading. Finally, we asked whether exocyst mutants would produce cargo distribution defects in other regions of the neuron. RNAi-mediated knockdown of Sec15 had no impact on the levels of Syt4-GFP in cell bodies or axons (**Fig. 4B-C**). This finding indicates a local role for Exocyst in maintaining proper EV cargo levels at synaptic terminals.

**Figure 4.**
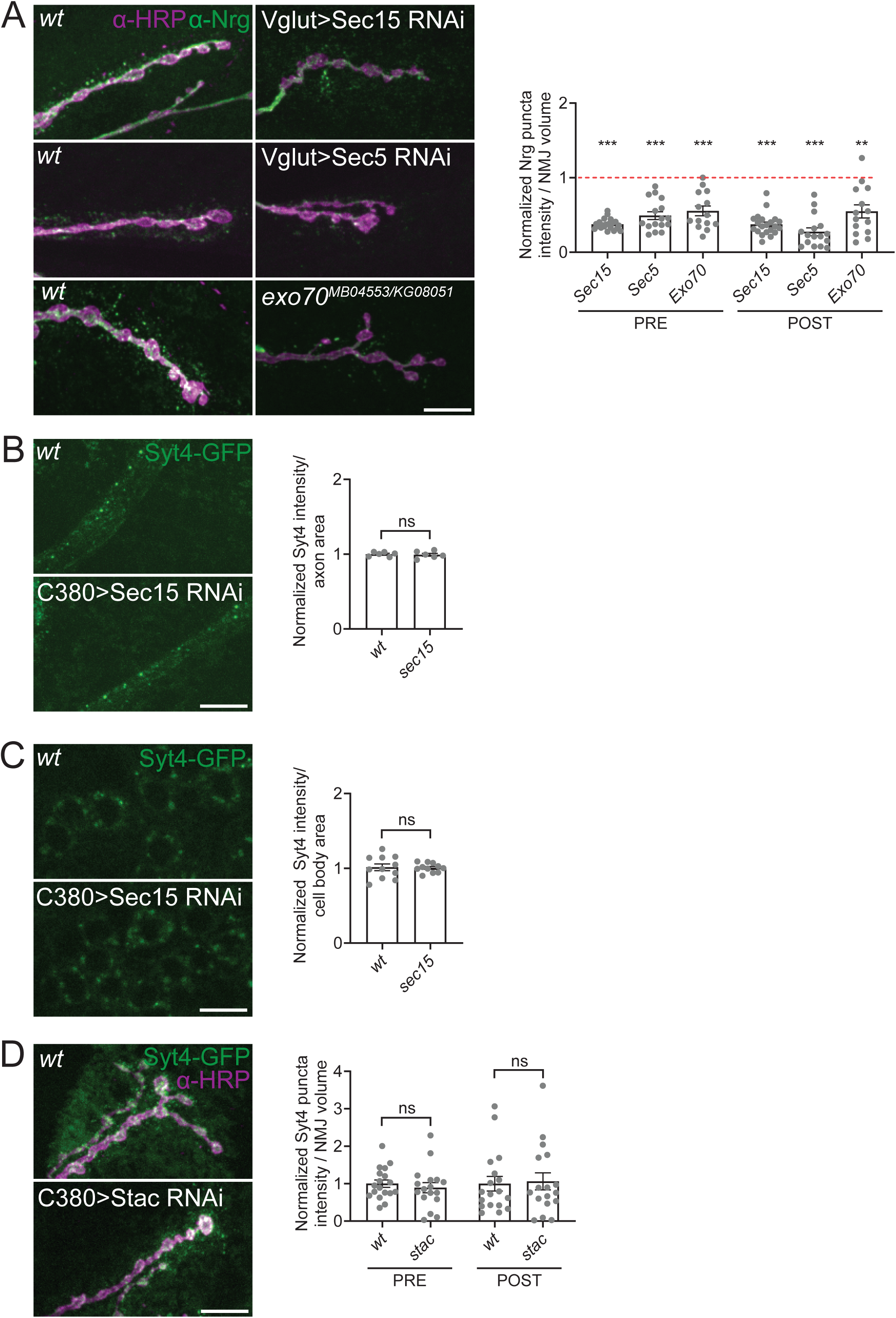
Tethering effector proteins indicate a role for Rab11 in recycling flux rather than MVB fusion. (A-C) Loss of components of the exocyst complex (*sec15, sec5,* and *exo70*) leads to a depletion of Nrg from NMJs, with loss of exocyst complex component *sec15* showing no change in levels of Syt4GFP in axons or cell bodies. **(A)** Representative images of Nrg at muscle 6/7 NMJs and quantification of pre-and postsynaptic Nrg puncta intensity. **(B)** Representative images of Syt4-GFP in distal axons and quantification of mean intensity. **(C)** Representative images of Syt4-GFP in cell bodies and quantification of mean intensity. **(D)** Loss of *stac* leads to no change in Syt4-GFP levels at the NMJ. Representative images of Syt4-GFP at muscle 6/7 NMJs and quantification of pre- and postsynaptic Syt4-GFP puncta intensity. Data is represented as mean ± SEM; n is depicted by individual gray dots in the graphs and represents NMJs **(A,D)**, axon average per animal **(B)**, or brains **(C)**. Cell body and axon intensity measurements were normalized to area of compartment (as noted in graphs), NMJ intensity measurements were normalized to presynaptic volume; all measurements were further normalized to the mean of their respective controls. All scale bars are 10 µm. See **Tables S1** and **S2** for detailed genotypes, sample sizes, and statistical analyses. wt, wild-type. α-HRP was used to label neuronal membranes.

We next investigated Staccato (Stac, mammalian Munc13-4), a calcium-binding protein involved in vesicle tethering and fusion. Stac binds Rab11 in multiple cell types and promotes Rab11 vesicle trafficking in a Ca^2+^-dependent manner in neutrophils (Gillingham et al., 2014; Johnson et al., 2016). In cancer cells, Munc13-4 facilitates Ca^2+^-dependent interactions or fusion between recycling endosomes and MVBs, enhancing exosome release (Messenger et al., 2018). In *Drosophila*, Stac localizes to MVBs and EVs within developing tracheal tubes (Camelo et al., 2022). After validating an RNAi line against Stac, which effectively depleted UAS-GFP-StacA at the NMJ (**Fig S2D**), we examined Syt4-GFP levels in motor neurons expressing UAS-Stac-RNAi. No significant changes were observed pre- or postsynaptically (**Fig. 4D**), suggesting that Stac is not critical for EV cargo trafficking at the NMJ, consistent with its low mRNA expression in motor neurons (Jetti et al., 2023).

Collectively, these findings highlight the roles of tethers and fusion-promoting factors in regulating EV cargo flux at the NMJ, and support a model in which Rab11 primarily drives recycling flux rather than directly mediating MVB fusion.

### Selective involvement of Rab11-interacting sorting proteins in EV trafficking

We next explored the roles of several proteins with sorting functions identified as putative Rab11-interactors in a proteomics screen from *Drosophila* S2 cells (Gillingham et al., 2014). One candidate was Receptor Mediated Endocytosis 8 (Rme-8, or DNAJC13 in mammals), which, although not yet validated as a Rab11-effector, is crucial for endosomal sorting and selective cargo endocytosis (Chang et al., 2004; Girard et al., 2005; Girard & McPherson, 2008; Zhang et al., 2001). Rme-8 also interacts with the Retromer complex, which opposes Rab11 in several processes, including EV trafficking (Freeman et al., 2014; Popoff et al., 2009; Walsh et al., 2021), making it an interesting putative effector to investigate.

Rme-8 is essential for viability in *Drosophila*, and its neuronal knockdown causes lethality in adult flies (Gomez-Lamarca et al., 2015; Raut et al., 2017). Despite these established roles, we found that neuronal knockdown of Rme-8 did not affect Nrg levels at either pre- or postsynaptic sites (**Fig. 5A**). This indicates that Rme-8 does not act in the same pathway as Rab11 or in opposition to Rab11 in regulating EV cargo trafficking. These findings suggest that, despite its known role in endosomal recycling and its identification as a putative Rab11 interactor, Rme-8 does not influence EV trafficking at the NMJ, and reinforce the specificity of the phenotypes observed for other Rab11-associated trafficking factors.

**Figure 5.**
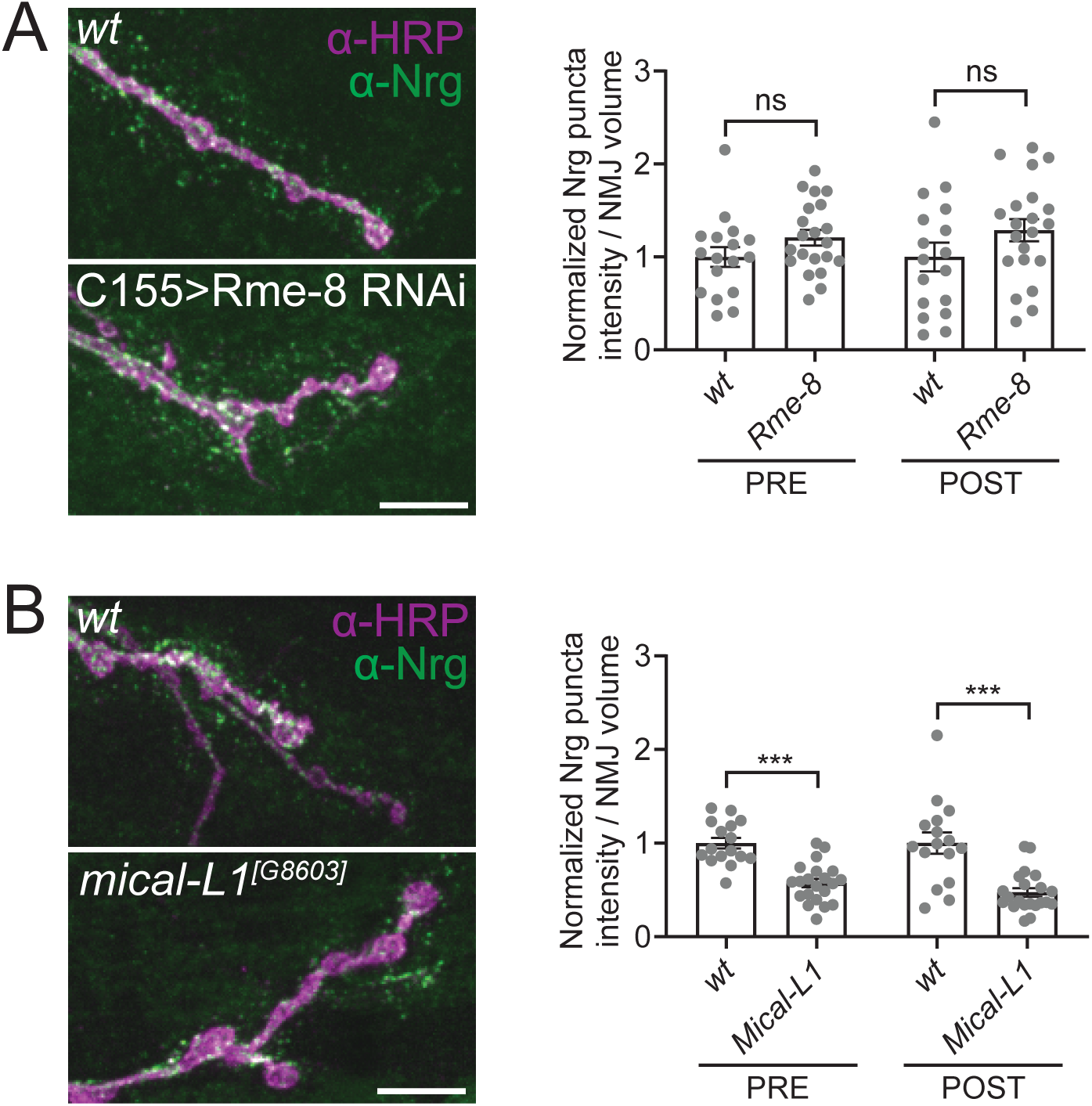
Select recycling regulators promote EV cargo trafficking. **(A)** Loss of *rme-8* leads to no change in Nrg levels at NMJs. Representative images of Nrg at muscle 6/7 NMJs and quantification of pre-and postsynaptic Nrg puncta intensity. **(B)** Loss of *mical-L1* leads to depletion of Nrg at NMJs. Representative images of Nrg at muscle 6/7 NMJs and quantification of pre- and postsynaptic Nrg puncta intensity. Data is represented as mean ± SEM; n is depicted by individual gray dots in the graphs and represents NMJs. NMJ intensity measurements were normalized to presynaptic volume; all measurements were further normalized to the mean of their respective controls. All scale bars are 10 µm. See **Tables S1** and **S2** for detailed genotypes, sample sizes, and statistical analyses. wt, wild-type. α-HRP was used to label neuronal membranes.

Next, we investigated Mical-like (Mical-L1), a protein with diverse roles in endosomal recycling, making it a compelling candidate for regulating EV trafficking. Although no direct interaction between Rab11 and Mical-L1 has been documented, Mical-L1 functions downstream of several Rab GTPases, including Rab11, in mammalian cells, where it promotes tubule formation. Through its association with retromer, Mical-L1 is hypothesized to facilitate the scission of these tubules (Giridharan et al., 2013; Naslavsky & Caplan, 2018). Furthermore, a recent study revealed a physical interaction between Mical-L1 and the F-BAR/SH3 membrane remodeling protein FCHSD2, showing that Mical-L1 recruits FCHSD2 to endosomes, enabling tubule fission and receptor recycling (Frisby et al., 2024). Notably, in *Drosophila*, mutants of the FCHSD2 homolog Nervous wreck (Nwk) display local depletion of EV cargoes at synapses, similar to *rab11* mutants (Blanchette et al., 2022). Consistent with a role in synaptic recycling, we found that a previously described P-element insertion into the coding region of Mical-L1, likely resulting in a null allele (Nahm et al., 2016), led to decreased Nrg levels at both pre- and postsynaptic sites (**Fig. 5B**). Overall, our results indicate that only select Rab11 interactors with sorting functions, such as Mical-L1, play a role in promoting EV trafficking, perhaps in cooperation with membrane remodeling machinery such as Nwk.

### Distinct PI(4)P regulators bidirectionally control EV trafficking

Enzymes controlling the phospholipid composition of membranes are frequently regulated by Rab GTPases (Pfeffer, 2017). To investigate the role of phosphatidylinositol synthesis in EV trafficking, we first tested the phosphatidylinositol-4 kinase Four wheel drive (Fwd, PI4KIIIβ in mammals) which localizes to Golgi membranes and Golgi-derived vesicles, where it synthesizes PI(4)P (Audhya et al., 2000; Godi et al., 1999; Polevoy et al., 2009; Sciorra et al., 2005). Fwd binds to Rab11 independently of both its own kinase activity and Rab11’s GTP-binding state, positioning it upstream of Rab11 (Burke et al., 2014; de Graaf et al., 2004; Polevoy et al., 2009). To assess the role of *fwd* in EV trafficking, we measured EV cargo levels in larvae carrying a trans-heterozygous combination of a *fwd* mutant and a deficiency (Polevoy et al., 2009). We observed a significant increase in both Nrg (**Fig. 6A**) and Syt4-GFP (**Fig. 6B**) levels at both pre- and postsynaptic sites. We also observed a mild increase in the levels of Syt4-GFP in motor neuron cell bodies (**Fig. 6C**) and axons (**Fig. 6D**) in *fwd* mutants.

**Figure 6.**
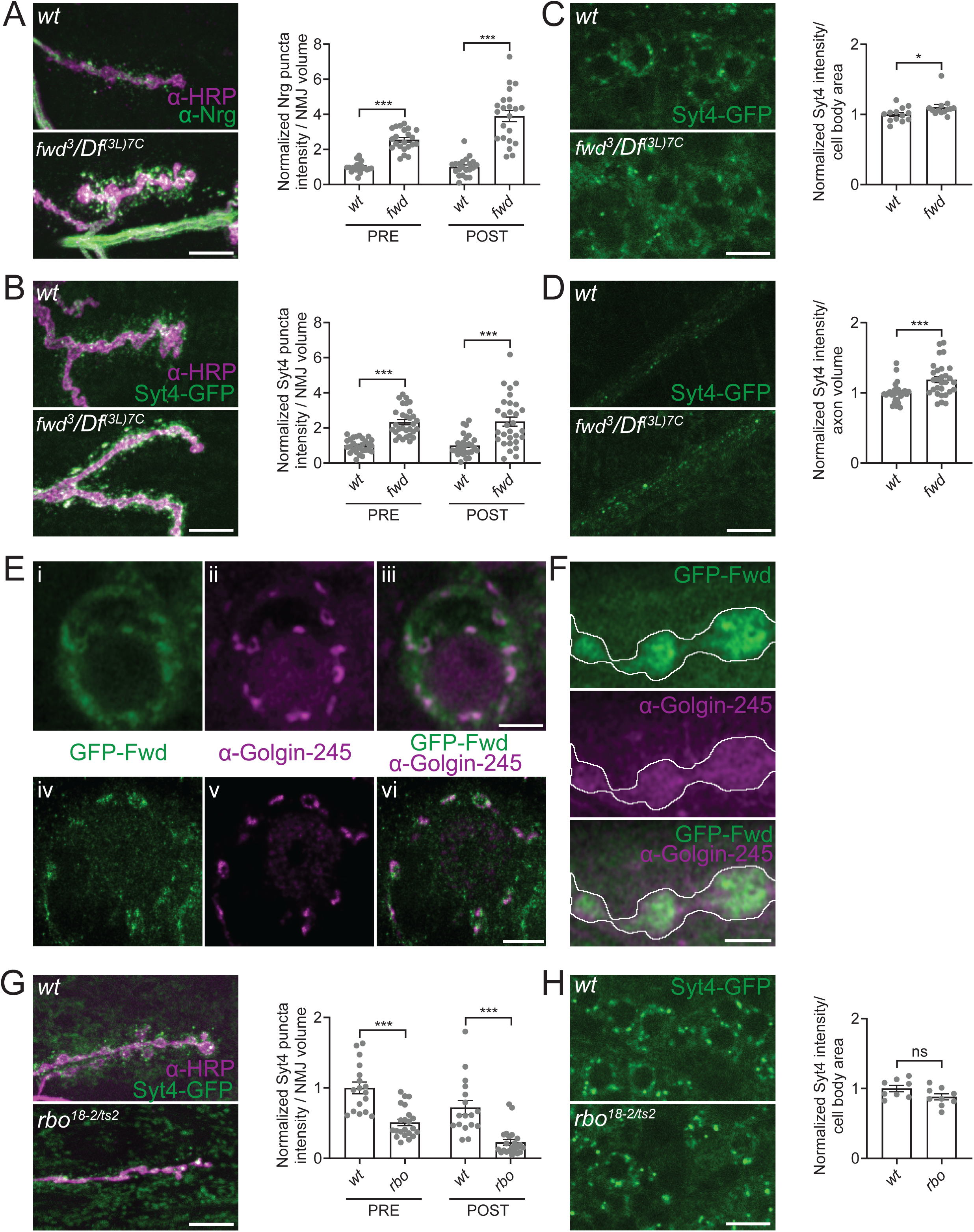
Two PI(4) kinases differentially regulate EV trafficking. **(A)** Loss of *fwd* leads to an increase in Nrg at NMJs. Representative images of Nrg at muscle 6/7 NMJs and quantification of pre- and postsynaptic Nrg puncta intensity. **(B-D)** Loss of *fwd* leads to an increase in Syt4-GFP levels at NMJs, cell bodies, and axons. **(B)** Representative images of Syt4-GFP at muscle 4 NMJs and quantification of pre-and postsynaptic Syt4-GFP puncta intensity. **(C)** Representative images of Syt4-GFP in cell bodies and quantification of mean intensity. **(D)** Representative images of Syt4-GFP in distal axons and quantification of mean intensity. **(E)** Single Z slices from stacks taken with Airyscan (i,ii,iii) or 3D STED (iv, v, vi) microscopy of a motor neuron cell body expressing GFP-Fwd and labeled with the trans-Golgi marker Golgin-245. **(F)** Single Z slices from stack taken with Airyscan of an NMJ expressing GFP-Fwd and labeled with the trans Golgi marker Golgin-245. **(G-H)** Loss of *rbo* leads to depletion of Syt4-GFP from NMJs with no change in cell bodies. **(G)** Representative images of Syt4-GFP at muscle 6/7 NMJs and quantification of pre- and postsynaptic Syt4-GFP puncta intensity. **(H)** Representative images of Syt4-GFP in cell bodies and quantification of mean intensity. Data is represented as mean ± SEM; n is depicted by individual gray dots in the graphs and represents NMJs **(A,B,G)**, axons **(D)**, or brains **(C,H)**. Cell body and axon intensity measurements were normalized to area or volume of compartment (as noted in graphs), NMJ intensity measurements were normalized to presynaptic volume; all measurements were further normalized to the mean of their respective controls. Scale bars are 10 µm **(A, B, C, D, G, H)** or 2 µm **(E, F)**. See **Tables S1** and **S2** for detailed genotypes, sample sizes, and statistical analyses. wt, wild-type. α-HRP was used to label neuronal membranes.

These results mirror our previous findings in *vps35* mutants, which disrupt the Retromer complex (Walsh et al., 2021). Retromer retrieves cargoes from endosomes for transport to either the Golgi or plasma membrane (Chen et al., 2019). The similar phenotypes in *fwd* and *vps35* mutants, combined with Fwd’s known Golgi localization, raise a possibility role for Golgi in EV trafficking. To explore potential Golgi involvement, we examined Fwd localization. In motor neuron cell bodies, UAS-GFP-Fwd showed strong colocalization (by both Airyscan and STED microscopy) with the trans-Golgi marker Golgin-245 (**Fig 6E**). At presynaptic NMJ terminals, UAS-Fwd-GFP was detectable by Airyscan microscopy in a reticular pattern, though it was too dim for STED analysis **(Fig. 6F**). Importantly, we did not observe specific presynaptic Golgin-245 staining at terminals. (**Fig. 6F**). These results raise the possibility that Fwd may have presynaptic functions independent of conventional Golgi.

We further examined PI(4)P synthesis by investigating Rolling blackout (Rbo), a component of the PI4KIIIα complex, which converts PI to PI(4)P, likely at the plasma membrane (Nakatsu et al., 2012). Although the PI4KIIIα complex has not been previously linked to Rab11 or studied in the context of EV trafficking, all five components were identified as Rab11 interactors in S2 cells (Gillingham et al., 2014). Since *rbo* null mutants are lethal, we analyzed a heterozygous combination of a truncated version of the protein (*rbo^18-2^*) and a temperature-sensitive allele (*rbo^ts2^*). This allelic combination is viable in third instar larvae and exhibits membrane trafficking defects at the NMJ (Huang et al., 2004, 2006; Liu et al., 2018).

We observed a significant decrease in Syt4-GFP levels at both pre- and postsynaptic sites in *rbo^ts2^/rbo^18-2^* mutants raised at the permissive temperature for 48 hours followed by the restrictive temperature for 96 hours (**Fig. 6G**). Further, when we examined Syt4-GFP levels in the cell bodies of *rbo^ts2^/rbo^18-2^* mutants, we saw no change compared to controls (**Fig. 6H**), suggesting that Rbo exerts its role specifically at synapses. These findings suggest that the PI4KIIIα complex may regulate lipid composition to promote EV trafficking at the synapse, and supports its possible function as a novel Rab11 effector.

## Discussion

### Rab11 Regulates EV Cargo Trafficking and Localization in Neurons

In this study, we demonstrate that Rab11 acts as a key regulator of EV cargo localization in neurons. We found that in *rab11* loss-of-function mutants, EV cargoes are redistributed from motor neuron synapses to axons and cell bodies. Our live imaging analysis provides mechanistic insight into this redistribution, by revealing an increased number of stationary EV cargo-containing vesicles in axons of *rab11* hypomorphs without changes in cargo directionality or transport rates. One hypothesis explaining presynaptic depletion of EV cargoes is that *rab11* mutants impair EV cargo entry into the NMJ. However, this seems unlikely given that we see cargo increases in both proximal and distal axons. Instead, based on our previous findings showing that disruptions in endocytic recycling pathways lead to Dynein/Dynactin-mediated retrograde transport of EV cargoes (Blanchette et al., 2022), we propose that *rab11* mutants locally disrupt trafficking events at synapses, leading to cargo accumulation in membrane compartments that are then routed for retrograde transport. This model explains both the synaptic depletion and the axonal accumulation we observe. Notably, we observed cargo-specific differences in cell body accumulation, where overexpressed APP-GFP accumulated while endogenous Syt4-GFP levels remained unchanged. This likely reflects either differences in degradative capacity for the overexpressed APP-GFP, or distinct trafficking routes for each of these cargoes. Overall, our findings suggest that Rab11’s primary role is to properly maintain EV-cargo-containing compartments at synaptic terminals, rather than in long-distance transport. This led us to systematically examine how Rab11’s diverse effectors contribute to synaptic EV cargo trafficking.

### Rab11 Maintains EV Cargo Pools Through Recycling Flux

Previous studies suggested that Rab11, MyoV and Exocyst facilitate MVB-plasma membrane fusion based on postsynaptic EV cargo depletion in *rab11* mutants (Koles et al., 2012; Messenger et al., 2018; Savina et al., 2005; Kang et al., 2024; Bai et al., 2022). However, this fusion model predicts presynaptic cargo accumulation coupled with postsynaptic depletion (as seen in Hsp90 mutants where stalled MVBs accumulate near the plasma membrane (Lauwers et al., 2018)), while a recycling flux model predicts overall cargo depletion at both sites. Since we previously observed the latter pattern in *rab11* mutants (Walsh et al., 2021), and disruption of Rab11 effectors like MyoV and Exocyst produces similar cargo depletion rather than the Hsp90-like phenotype, our results support a model where Rab11 maintains EV cargo pools through recycling flux rather than directly mediating MVB fusion. However, we cannot fully exclude a secondary role in MVB fusion occurring alongside cargo flux regulation.

Our results also reveal unexpectedly opposing functions for Rab11’s motor protein effectors. MyoV promotes EV trafficking while Nuf restricts it, suggesting that Rab11 orchestrates bidirectional control of EV cargo distribution. Notably, *nuf* mutants show cargo accumulation both pre- and postsynaptically, unlike the Hsp90 phenotype where fusion is blocked causing only presynaptic accumulation. This indicates that Nuf regulates flux in the opposite direction from MyoV rather than having a distinct function in fusion. MyoV likely promotes local transport of recycling endosomes to the plasma membrane via actin-based movement, maintaining EV cargo availability at terminals (Ruiz-Cañada & Budnik, 2006). Conversely, Nuf may facilitate cargo removal through its interactions with dynein motor complexes, enabling retrograde transport of EV cargo-containing compartments from synaptic terminals (Blanchette et al., 2022). Interestingly, the increase in both pre- and postsynaptic EV cargoes upon Nuf knockdown contrasts with the strictly presynaptic increase seen with dominant negative Dynactin (Blanchette et al., 2022). Since both Nuf and Dynactin regulate Dynein-mediated transport but produce opposite phenotypes, this suggests they control different stages in the maturation of EV precursor compartments. Nuf disruption may arrest compartments at an earlier, EV-competent stage, while Dynactin disruption may arrest them later, after the compartments have matured beyond the stage where they can contribute to EV release. Overall, our results indicate that Rab11 controls a multistep and bidirectional process that allows precise control of endosomal EV cargo pools available for release.

### EV cargo sorting on endosomes

Efficient trafficking in the EV pathway requires that cargoes navigate through multiple compartments while avoiding alternative pathways. Our previous and new data suggest that proteins such as Retromer (Walsh et al., 2021) and Mical-L1 play roles in sorting EV cargoes into release-competent compartments. The specificity of this EV cargo sorting function is underlined by our finding that loss of the ubiquitous sorting factor *rme-8* does not cause detectable EV cargo sorting defects, despite its expression in motor neurons (Jetti et al., 2023) and its requirement for central nervous system structure (Lacin et al., 2024).

We previously found a prominent role for the F-BAR/SH3 protein Nwk, along with other canonical endocytic proteins, in maintaining an available pool of EV cargoes locally at the NMJ (Blanchette et al., 2022), similar to Rab11. Interestingly, Mical-L1 mutants show several additional phenotypes resembling those of *nwk* mutants, including synaptic overgrowth and misrouting of cargoes to Spinster-positive degradative compartments, in addition to depleting EV cargoes from synapses (Blanchette et al., 2022; Nahm et al., 2016). Together with recent evidence for a physical interaction between MICAL-L1 and the Nwk homolog FCHSD2 in mammalian cells, our results suggest that Mical-L1 and Nwk may work together to promote EV cargo trafficking (Frisby et al., 2024). These data add to the mounting evidence that Nwk/FCHSD2 and associated membrane remodeling proteins may function on intracellular compartments, beyond their canonical role in regulating force-generating actin assembly at the plasma membrane during the internalization step of endocytosis (Almeida-Souza et al., 2018; Del Signore et al., 2021; Xiao et al., 2018). FCHSD2 is recruited by MICAL-L1 to endosomes, where it regulates actin assembly and is required for recycling of Transferrin, major histocompatibility complex class I (MHC I) receptors, and receptor tyrosine kinases (Frisby et al., 2024; Xiao & Schmid, 2020). Further, *Drosophila* Nwk interacts with the endosomal sorting protein SNX16 to control trafficking of signaling receptors through tubulated endosomal intermediates (Rodal et al., 2011; Wang et al., 2019). Whether the EV trafficking defects we observe result from Nwk’s canonical endocytic functions versus a novel role in endosomal cargo sorting (together with Mical-L1) will likely require the identification of separation-of-function mutants that distinguish these activities.

### PI(4)P as a Potential Sorting Signal for EV Cargoes

Our findings show that two different PI(4)P-synthesizing proteins influence EV trafficking in opposite directions, positioning PI(4)P as an important sorting signal for EV cargoes. Rbo, which functions as a component of the PI4KIIIα complex to generate PI(4)P at the plasma membrane (Chung et al., 2015; Polevoy et al., 2009), promotes EV trafficking. Fwd, which is thought to generate PI(4)P at the Golgi (Polevoy et al., 2009), restricts EV cargo flux. While future work should investigate whether these phenotypes depend on the catalytic activity of Rbo and Fwd or arise from other functions of these proteins, our results suggest that distinct PI(4)P pools may act at different locations to regulate EV cargo traffic. The known functions of Rbo and Fwd provide some clues to the nature of these different pools.

Rbo was originally characterized as essential for activity-dependent bulk endocytosis (ADBE), which is a physiologically important mode of synaptic membrane retrieval after intense stimulation, and generates large cisternae that can be resolved into various endocytic and endosomal compartments (Vijayakrishnan et al., 2009, Korber et al., 2012; Wenzel et al., 2012; Arpino et al., 2022; Chanaday et al., 2019; Clayton et al., 2008; Ivanova and Cousin, 2022). Interestingly, Rab11 also regulates ADBE, and co-fractionates with ADBE-generated endosomes (Kokotos et al., 2018). Our finding that *rbo* mutants exhibit reduced EV cargo levels at synapses, similar to *rab11* mutants, suggests a novel mechanistic link between Rab11, PI4KIIIα, ADBE and EV trafficking, which is further supported by proteomics studies showing that all PI4KIIIα complex components were found in Rab11 pulldowns (Gillingham et al., 2014). This possible connection between ADBE and EV cargo trafficking suggests that the same Rab11/PI4KIIIα-dependent machinery that enables synaptic vesicle recycling during intense stimulation may simultaneously load EV precursor compartments, providing a novel mechanism for the poorly understood phenomenon of activity-dependent EV release (Faure et al., 2006; Fruhbeis et al., 2013; Lachenal et al., 2011; Olivero et al., 2021).

How then might Rbo-dependent PI(4)P synthesis contribute to EV cargo trafficking? ADBE requires PI(4,5)P2 microdomains at the plasma membrane (Li et al., 2020), suggesting that Rbo/PI4KIIIα might facilitate ADBE by providing PI(4)P as the rate-limiting substrate for PI(4,5)P2 synthesis (Trivedi et al., 2020). PI(4,5)P2 interacts with Exocyst to control Rab11 vesicle tethering (He et al., 2007), suggesting that Rab11 regulation of Rbo/PI4KIIIα could promote positive reinforcement of its own activities for sustained ADBE and EV cargo loading. Alternatively, since PI(4)P recruits Exocyst to MVBs for docking and release (Liu et al., 2023), it may similarly recruit other Rab11 effectors directly to endosomes to influence EV cargo sorting.

Fwd is best known as a Golgi-localized PI4KIIIβ that synthesizes PI(4)P and binds Rab11 independently of GTPase state (Burke et al., 2014; de Graaf et al., 2004; Audhya et al., 2000; Godi et al., 1999; Polevoy et al., 2009; Sciorra et al., 2005). We found that *fwd* mutants exhibit increased EV cargo levels at synapses, opposite to the *rab11* phenotype. The strong EV trafficking phenotype of a Golgi-associated protein was unexpected, as it is widely thought that the Golgi apparatus is absent from presynaptic terminals, and that EV cargo trafficking occurs through local endosomal recycling pathways (Patel et al., 2020; Blanchette et al., 2022).

This unexpected finding suggests three possible mechanisms for Fwd function at synapses. First, Fwd could exert Golgi-dependent effects from the cell body during EV cargo biosynthesis or processing, where loss of Fwd function leads to altered cargo modification or sorting that results in elevated steady-state levels. However, while we do observe a small but significant increase in EV cargo levels in cell bodies of *fwd* mutants, this increase is much smaller in magnitude than the dramatic accumulation at synapses, suggesting that cell body effects alone cannot fully explain the synaptic phenotype. Second, Fwd may have Golgi-independent functions at presynaptic terminals, consistent with our observation that UAS-GFP-Fwd localizes to synapses in a diffuse, reticular pattern despite the absence of canonical Golgi markers like Golgin-245. While this synaptic localization should be interpreted cautiously given the caveats of overexpression, it raises the possibility that Fwd functions on endosomal compartments independently of its canonical Golgi role. Finally, Fwd may act through local Golgi-like intermediates at presynaptic terminals that lack conventional markers or stacked morphology, similar to Golgi satellites/outposts or hybrid ER/Golgi compartments found in dendrites (Hertrich et al., 2024; Kennedy & Hanus, 2019; Mikhaylova et al., 2016; Ye et al., 2007). This putative local compartment could regulate EV cargo levels through post-translational modifications that promote cargo degradation, or by generating PI(4)P microdomains that recruit sorting machinery to divert EV cargoes away from release-competent compartments.

### Summary

Our findings reveal that Rab11 is essential for maintaining EV cargo localization within neurons, predominantly by regulating their trafficking rather than directly promoting release. We propose a model in which Rab11 coordinates motor proteins and phosphoinositides, particularly PI(4)P, to guide EV cargoes to release-competent compartments. By highlighting the complexities of Rab11 function, this study paves the way for future research to unravel the mechanisms of EV trafficking in neurons.

## Methods

### *Drosophila* strains and methods

*Drosophila* were grown at 25°C (or 29°C for RNAi experiments) on standard media. *rbo^ts^* crosses were grown at 25°C for 48 hours to allow adults time to lay eggs and then moved to 29°C. Crosses were density controlled, containing five virgin females and five males. Wandering 3^rd^ instar larvae were dissected in HL3.1 without calcium (Feng et al., 2004) on a Sylgard dish. Fillets were fixed in 4% paraformaldehyde in HL3.1 for 10 minutes while shaking at room temperature. Either male or female larvae were used for experiments, with each independent experiment using only one sex. See **Table S1** for detailed genotype information for each experiment and **Table S2** for *Drosophila* strain sources.

### Immunohistochemistry

Following fixation, larvae were washed four times for 5 minutes in PBS containing 0.2% TritonX-100 (0.2% PBX). Primary antibody details can be found in **Table S3**. Larvae were incubated with primary antibody in 0.2% PBX overnight at 4°C. Larvae were then washed 3-4 times for 5 minutes in 0.2% PBX and incubated with secondary antibodies or α-HRP antibodies diluted in 0.2% PBX (concentrations indicated in **Table S3**) for two hours at room temperature. For STED experiments, secondary labeling was done sequentially, with 5 10-minute washes in between secondaries in 0.2% PBX. Finally, larvae were washed 3-4 times for 5 minutes in 0.2% PBX. Larvae were then mounted in Vectashield (Vector Labs) or Abberior Mount Liquid anti-fade (Abberior) for spinning disk confocal imaging (mounting media kept consistent within each independent experiment), Abberior Mount Liquid anti-fade (Abberior) for Airyscan imaging or Prolong Diamond Antifade Mountant (ThermoFisher) for STED imaging.

### Image acquisition

All images were acquired at room temperature. Spinning disk confocal microscopy was conducted using Nikon Elements AR software on a Nikon Ni-E upright microscope a Yokogawa CSU-W1 spinning disk head, and an Andor iXon 897U EMCCD camera. For cell body and neuropil imaging of APP-GFP, we used a 40x (n.a. 0.75) oil immersion objective and 0.3 µm Z intervals, for cell body and axon imaging of Syt4-GFP, and fixed axon imaging of APP-GFP we used a 60x (n.a. 1.4) oil immersion objective and 0.3 µm Z intervals, and for muscle 6/7 NMJs in segments A2 and A3, we used a 60x (n.a. 1.4) oil immersion objective and 0.3 µm Z intervals. For axonal trafficking, time lapse images of axon bundles proximal to the ventral nerve cord were taken with 60x (n.a. 1.4) oil immersion objective. Nine Z slices with a 0.3µm step size and no acquisition delay between time points were collected per frame (frame rate 2.34-2.37 sec/frame). For Airyscan imaging, images were acquired at room temperature with Zen Black acquisition software on a Zeiss880 Fast Airyscan microscope in super resolution acquisition mode, using a 63x (n.a. 1.4) oil immersion objective and 0.18 µm Z intervals. STED images were acquired on an Abberior FACILITY line microscope with 60x (n.a. 1.3) silicone immersion objective, pulsed excitation lasers (561 nm and 640 nm), and a pulsed depletion laser (775 nm) to deplete all signals. Images were acquired at 60% 3D STED mode and pixel size was set to 30 nm.

### Image analysis

3D volumetric analysis of pre- and postsynaptic EV cargoes at NMJs was carried out using Volocity 6.0 software (Perkin Elmer) as previously described (Blanchette et al., 2022). For each NMJ image, only type Ib boutons were used for analysis, while axons and type Is arbors were cropped out. Presynaptic volume was determined through manual thresholding to the α-HRP signal (excluding objects smaller than 7µm^3^ and closing holes), with a 3.3µm dilation around this signal delineating the postsynaptic volume. Manual thresholding of the EV cargo signal was conducted to ensure measurements included only EV cargo signal above background muscle fluorescence. Postsynaptic objects smaller than 0.015µm^3^ were excluded. EV cargo sum intensity measurements were calculated within these pre- and postsynaptic volumes and normalized to the presynaptic volume.

Quantifications of EV cargoes in cell bodies and neuropil were conducted in FIJI (Schindelin et al., 2012). Individual groups of cell bodies (representing segmental repeats in the ventral ganglion) were manually traced in single confocal slices, and their mean intensity measured. A region within the neuropil was manually selected from a single slice through the center of the ventral ganglion. Mean EV cargo intensity within this region of neuropil was calculated.

Axonal EV cargo levels were measured in fixed larval fillets in either proximal axons (within 200 µm of the ventral ganglion) or distal axons (between the segmental axon bundle and synaptic boutons on muscle 4). Images were manually cropped to exclude everything aside from the axon segment to be measured. For data in **Fig. 1C** (APP-EGFP distal axons, sum slice projections) and **Fig. 4B** (Syt4-GFP distal axons, maximum intensity projections), the axon bundle was manually outlined using the α-HRP channel to select the region for measurement of mean APP-GFP intensity. For data in **Fig. 1D** (APP-EGFP proximal axons) sum slice projections were generated, and a background subtraction was done in FIJI with a rolling ball radius of 50 pixels with smoothing disabled. Thresholding was done on the HRP channel to generate a mask by manually selecting for each image the FIJI thresholding method with the best fit to the HRP signal of the axon bundle. APP-GFP mean intensity of the axon bundle was measured within this HRP mask. For data in **Fig. 1H** and **Fig. 6D (**Syt4-GFP or Nuf in distal axons), a background subtraction was done with a rolling ball radius of 50 pixels. Next a mask of the axon bundle in 3D was generated, using the α-HRP marker treated with a Gaussian blur. Finally, mean intensity of Syt4-GFP per axon volume was calculated by measuring the integrated density of Syt4-GFP within the 3D HRP mask and dividing this by the volume of the 3D HRP mask.

To estimate the distribution of EV cargoes between cell bodies, axons and NMJs, we modeled motor neuron cell bodies as spheres with a diameter of 10 µm, and axons as cylinders with a length of 1 mm and a diameter of 1 µm. We then measured the volume of NMJs from our data. Motor neuron dendrites were excluded from our estimates due to their relatively small volume compared to the cell body and axon (Kim et al., 2009). We measured mean intensity of EV cargoes in single confocal slices from 5 biological replicates in each of these regions. Assuming an axial point spread function of 1 µm, we multiplied this mean intensity by the estimated (cell body and axon) or calculated (NMJ) volume of each compartment to achieve a compartment integrated density.

To quantify APP-EGFP dynamics in live axons, maximum intensity projections of time course images were processed in FIJI to subtract background and adjust for XY drift using the StackReg plugin. Kymographs were generated from one to four axons per animal using the KymographBuilder plugin. Kymographs were blinded and the number of tracks were manually counted. The minimum track length measured was 3 µm, with most tracks greater than 5 µm. Velocity was measured by calculating the slope of the identified tracks. For each animal, the number of anterograde, retrograde, stationary or complex tracks were counted from one to four axons, and then divided by the total length of all measured axons from that animal.

### Statistics

All graphing and statistical analyses were conducted using GraphPad Prism. Datasets were first analyzed with the D’Agostino-Pearson normality test. Normally distributed datasets were analyzed with either an unpaired two-sided Student’s *t*-test (for datasets with two groups) or a one-way ANOVA with Tukey’s multiple comparisons (for datasets with more than two groups), while data that were not normally distributed were analyzed with either a two-sided Mann-Whitney U test (two groups) or a Kruskal-Wallis test with Dunn’s multiple comparisons (more than two groups). Error bars report ± SEM. Detailed information about genotypes, sample sizes, and statistical analyses performed for each dataset can be found in **Table S1**. * = P < 0.05; ** = P < 0.01; *** = P <0.001.

## Acknowledgements

We thank the Developmental Studies Hybridoma Bank created by the NICHD of the NIH, the Bloomington *Drosophila* Stock Center (Indiana University, Bloomington, IN, NIH P40OD018537), John Roche, Julie Brill, Anne Ephrussi, Akiko Satoh, and Stefan Luschnig for fly lines and antibodies, Iris Nava for technical assistance, and Julie Brill for helpful discussions. This work was supported by NINDS grant R01 NS103967 to A.A.R. and T32 MH019929 to A.L.S. and E.G., and by grant S10 OD034223 for the Abberior Facility Line STED microscope, housed in the Brandeis Light Microscopy Core Facility (RRID:SCR_025892).

---

**Figure S1.**
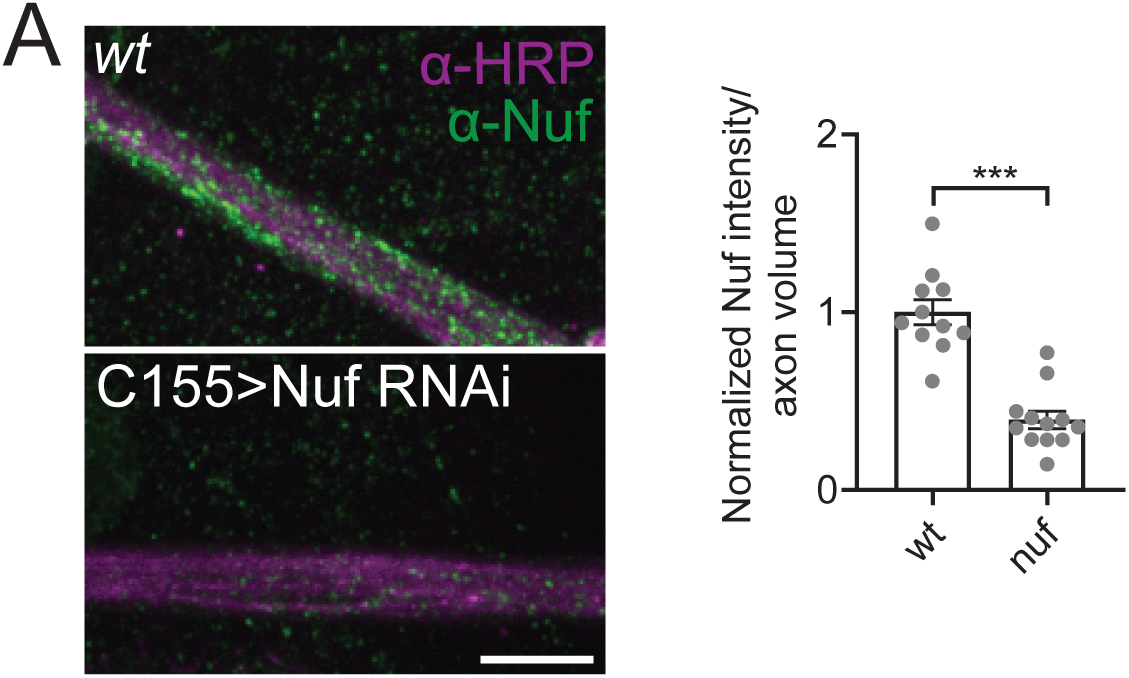
Validation of Nuf RNAi knockdown. **(A)** Representative images of α-Nuf antibody staining in axons demonstrating the neuronal Nuf RNAi knockdown effectively depletes Nuf levels in neurons and quantification of α-Nuf antibody levels in the axon bundle in control and upon neuronal Nuf RNAi knockdown. Data is represented as mean ± SEM; n is depicted by individual gray dots in the graph and represents axons. Axon intensity measurements were normalized to volume of compartment (as noted in graph); all measurements were further normalized to the mean of their respective controls. Scale bar is 10 µm. See **Tables S1** and **S2** for detailed genotypes, sample sizes, and statistical analyses. wt, wild-type. α-HRP was used to label neuronal membranes.

**Figure S2.**
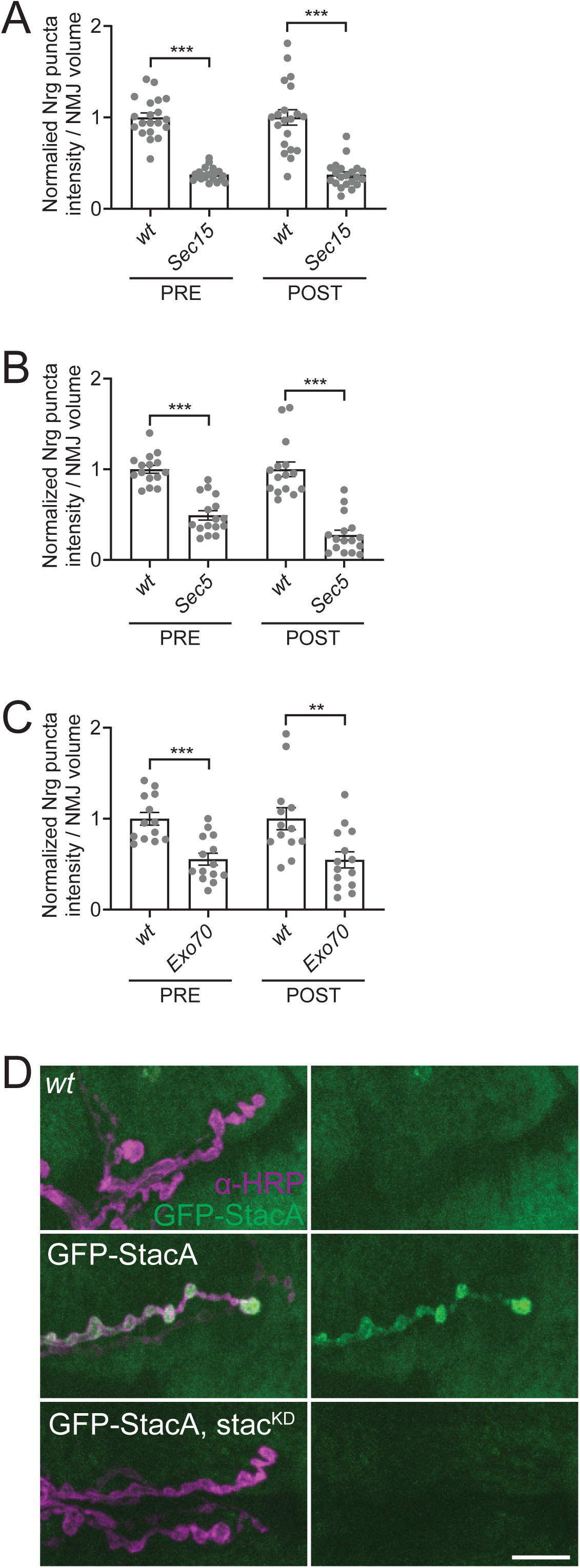
Tethering effector proteins support a role for Rab11 in recycling flux. Datasets pooled in Fig. 4AB shown individually with corresponding controls **(A)** Quantification of pre- and postsynaptic Nrg puncta intensity in control and Sec15 knockdown. **(B)** Quantification of pre- and postsynaptic Nrg puncta intensity in control and Sec5 knockdown. **(C)** Quantification of pre- and postsynaptic Nrg puncta intensity in control and *Exo70* mutants. **(D)** Representative images demonstrating that UAS-GFP-Stac expressed at the NMJ is depleted upon knockdown with UAS-Stac-RNAi. Data is represented as mean ± SEM; n is depicted by individual gray dots in the graphs and represents NMJs. NMJ intensity measurements were normalized to presynaptic volume; all measurements were further normalized to the mean of their respective controls. Scale bar is 10 µm. See **Table S1** and **S2** for detailed genotypes, sample sizes, and statistical analyses. wt, wild-type. α-HRP was used to label neuronal membranes.

**Table S1.**
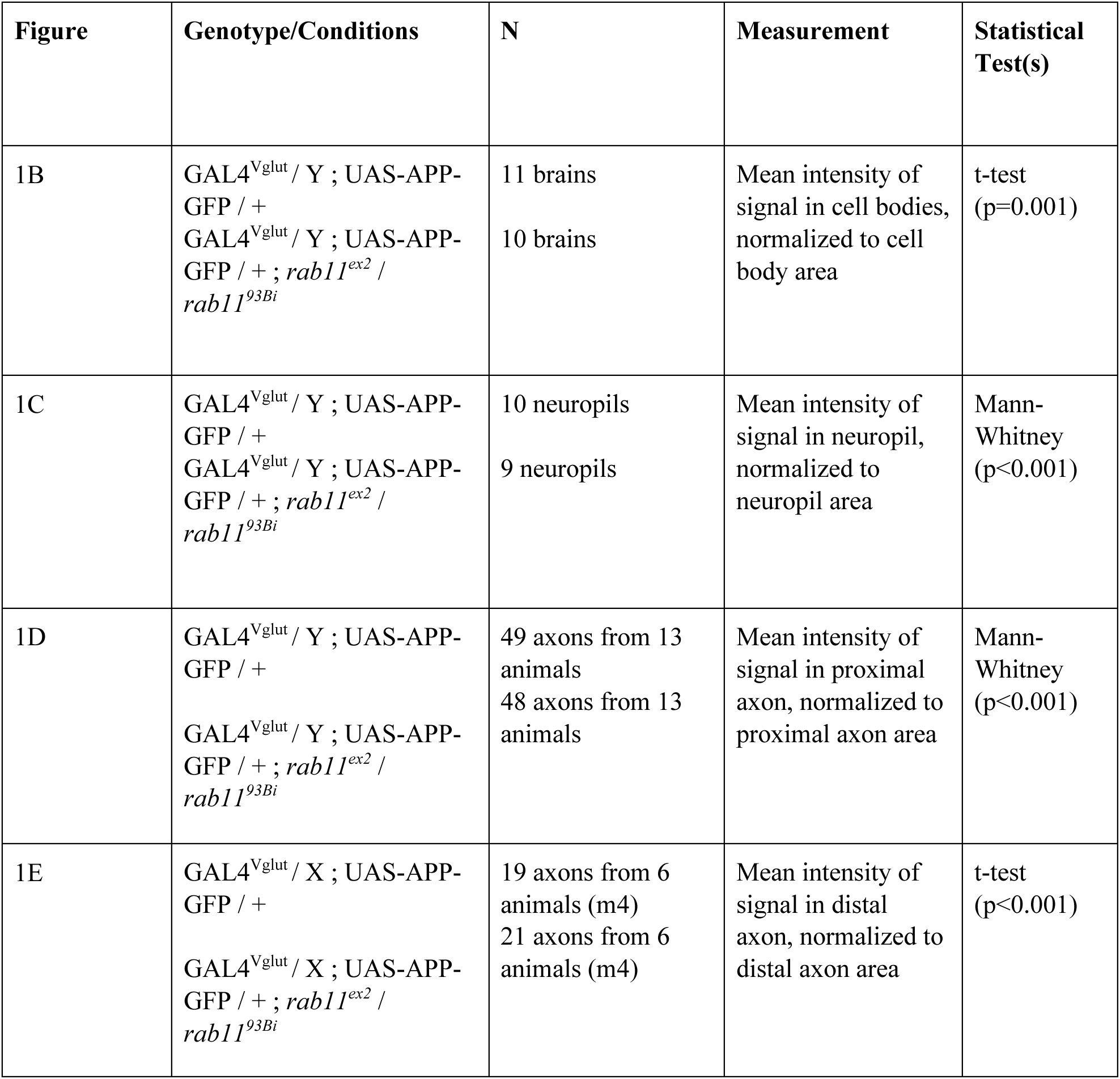
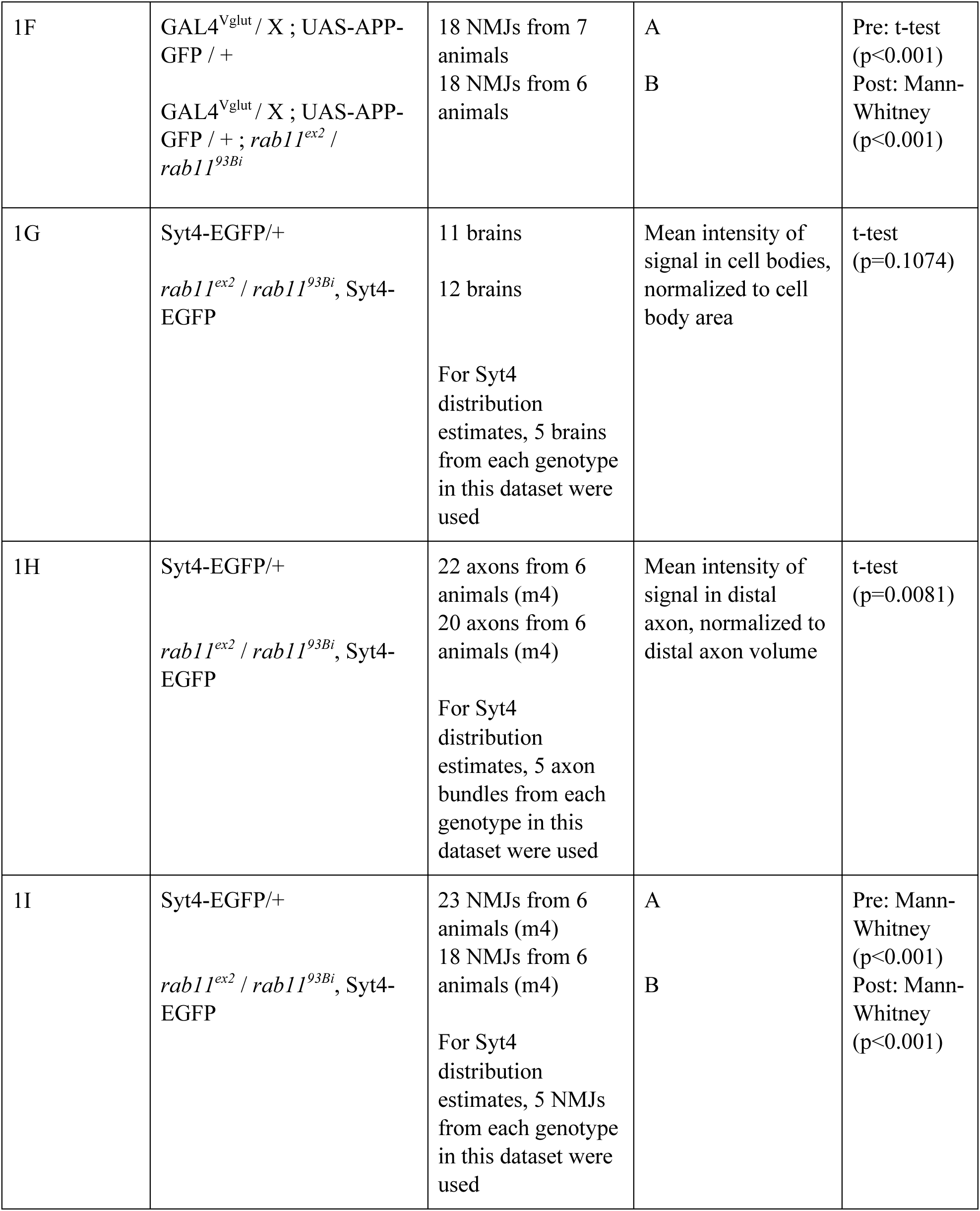
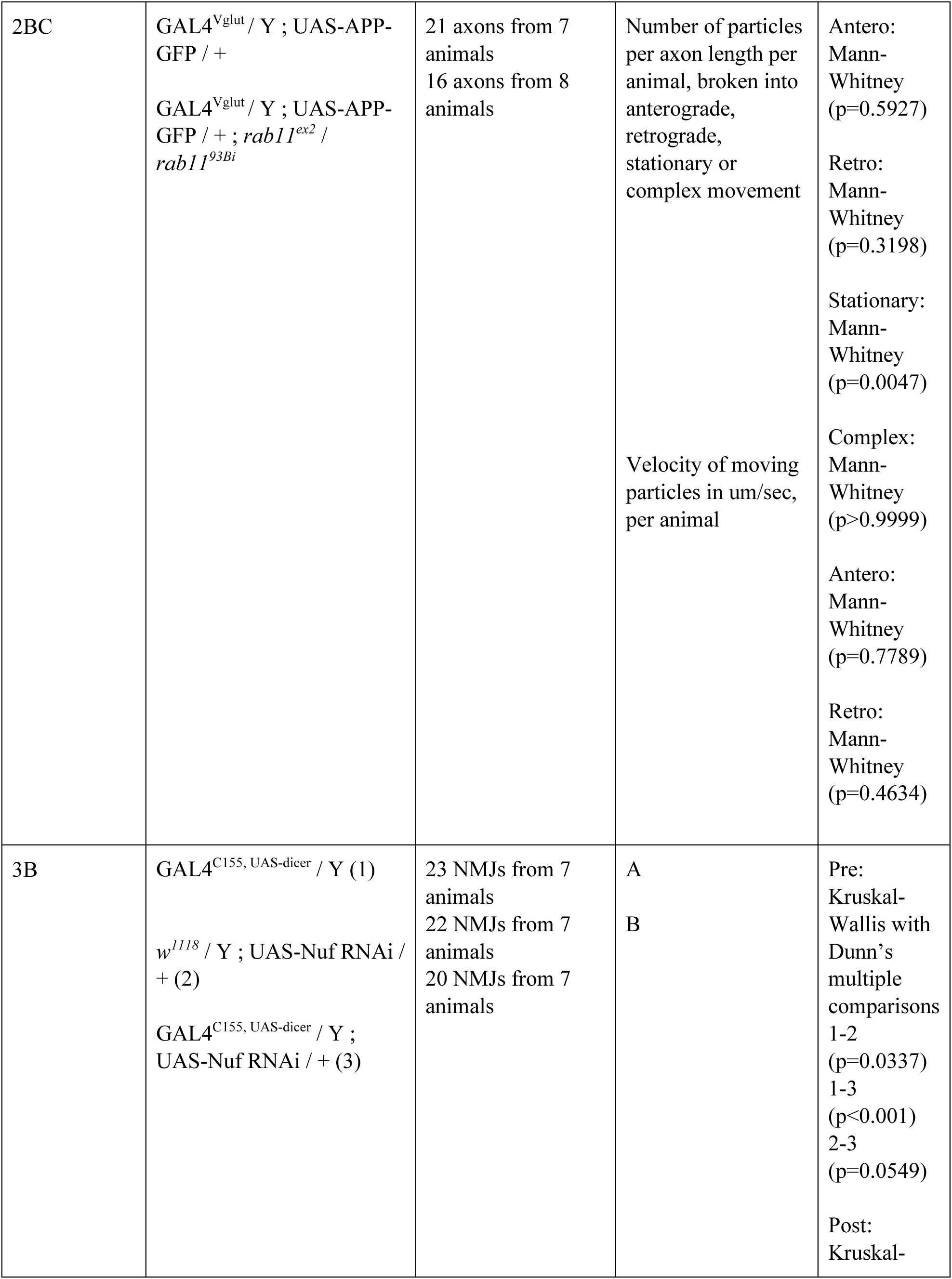
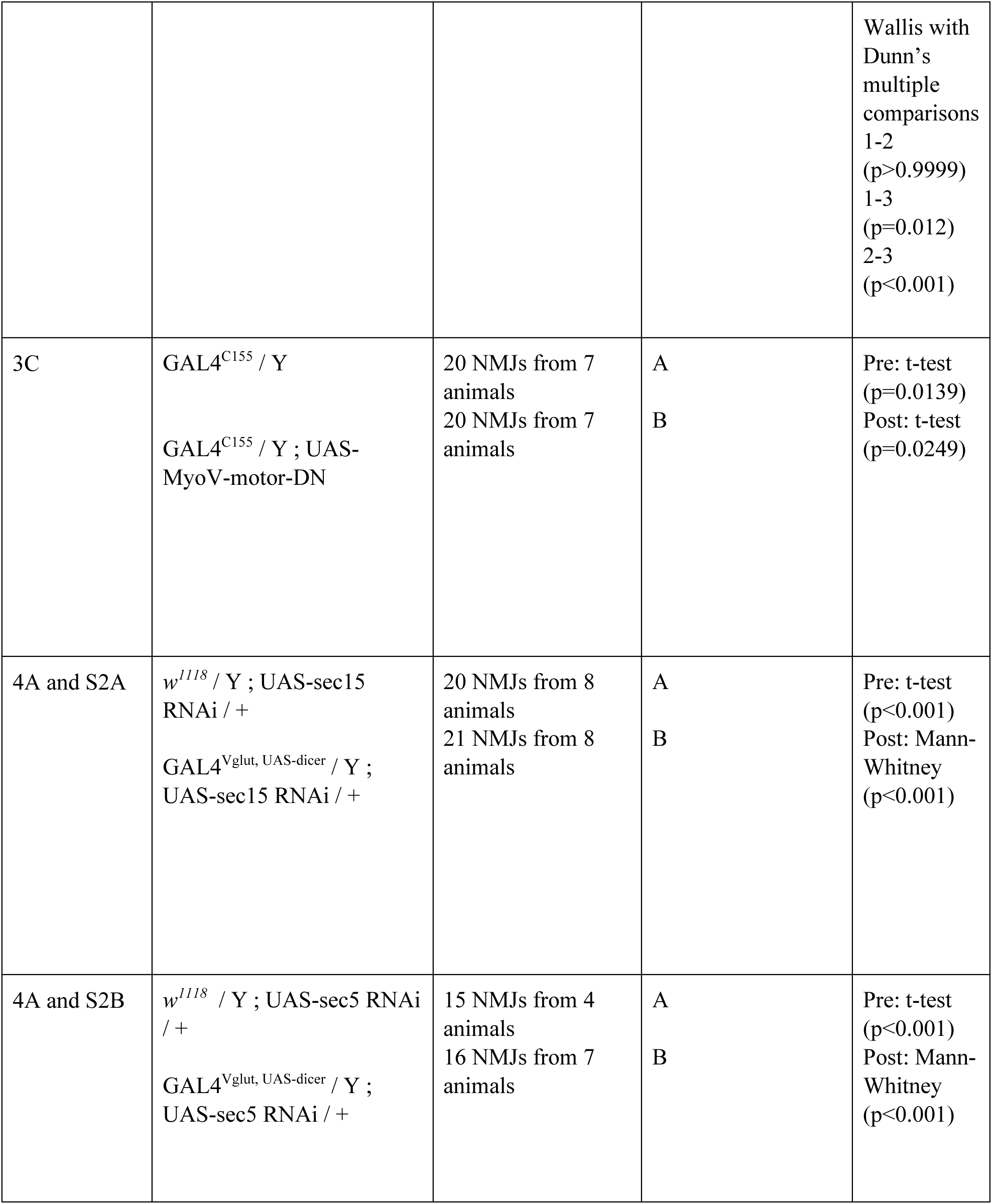
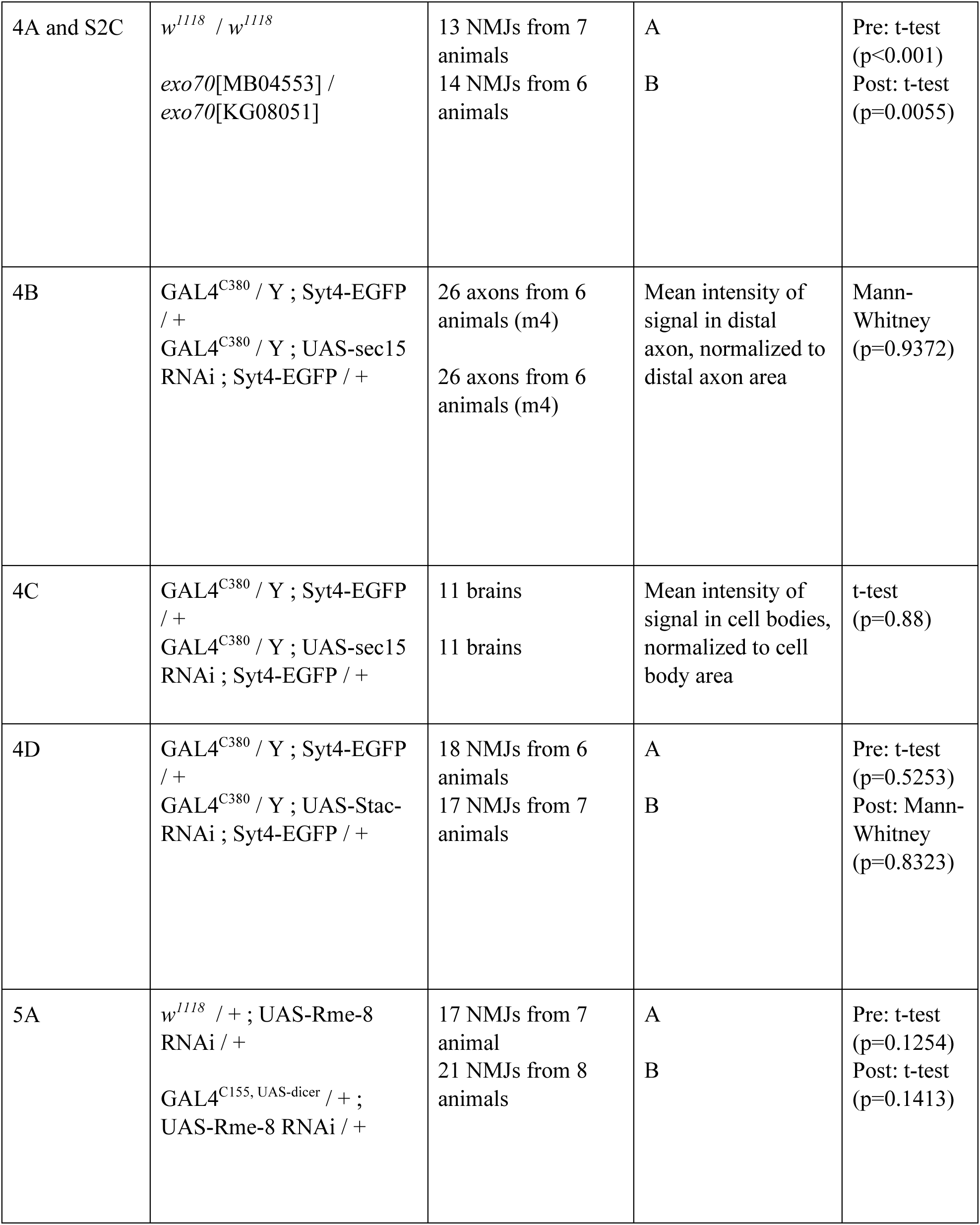
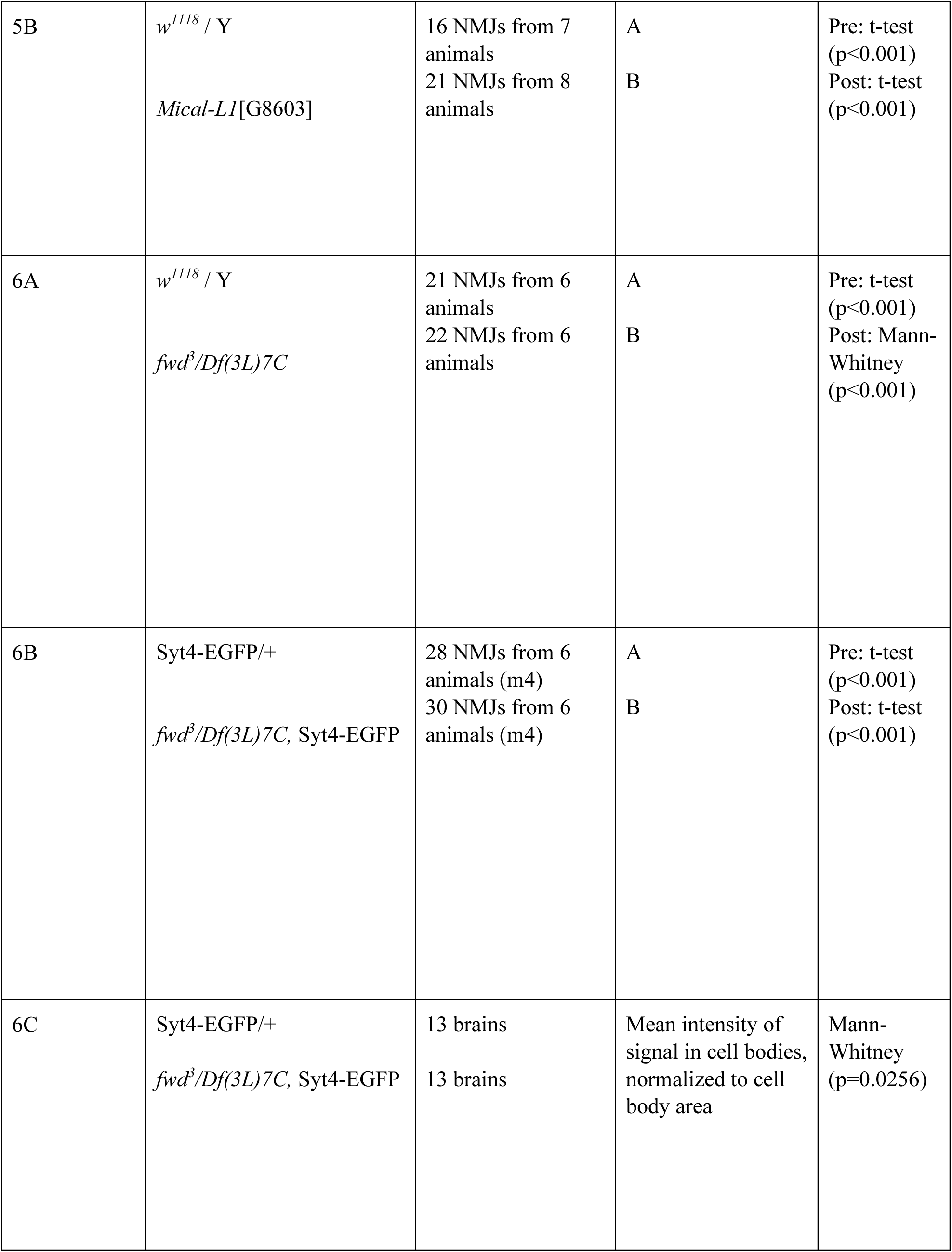
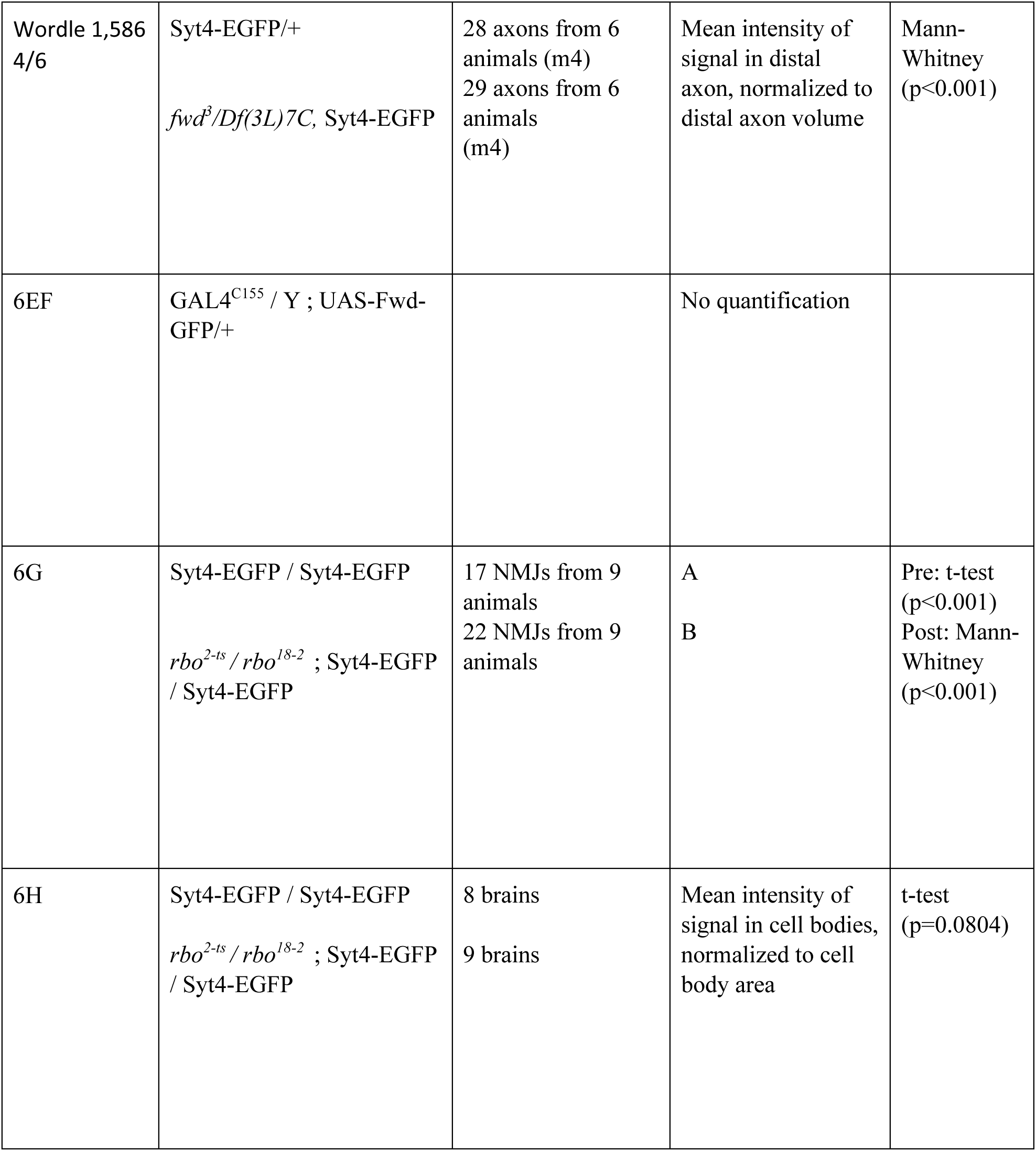
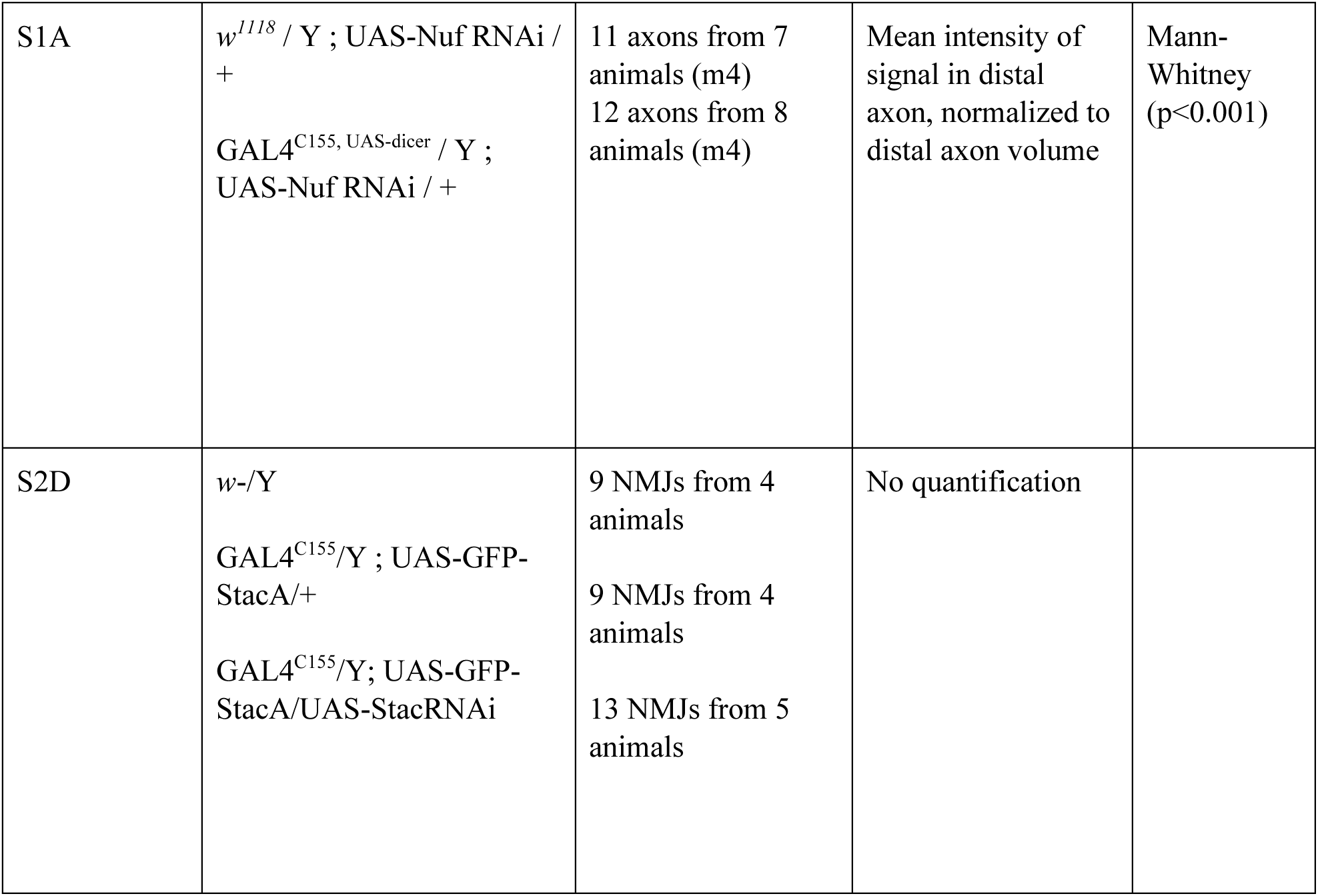
Statistics by dataset. Experiments were done at muscle 6/7 unless otherwise noted Presynaptic volume: α-HRP objects > 7μm^3^ Postsynaptic volume: 3 μm dilated from presynaptic volume A: Sum intensity of signal in thresholded objects in presynaptic volume, normalized to presynaptic volume B: Sum intensity of signal in thresholded objects in postsynaptic volume, normalized to presynaptic volume

**Table S2.**
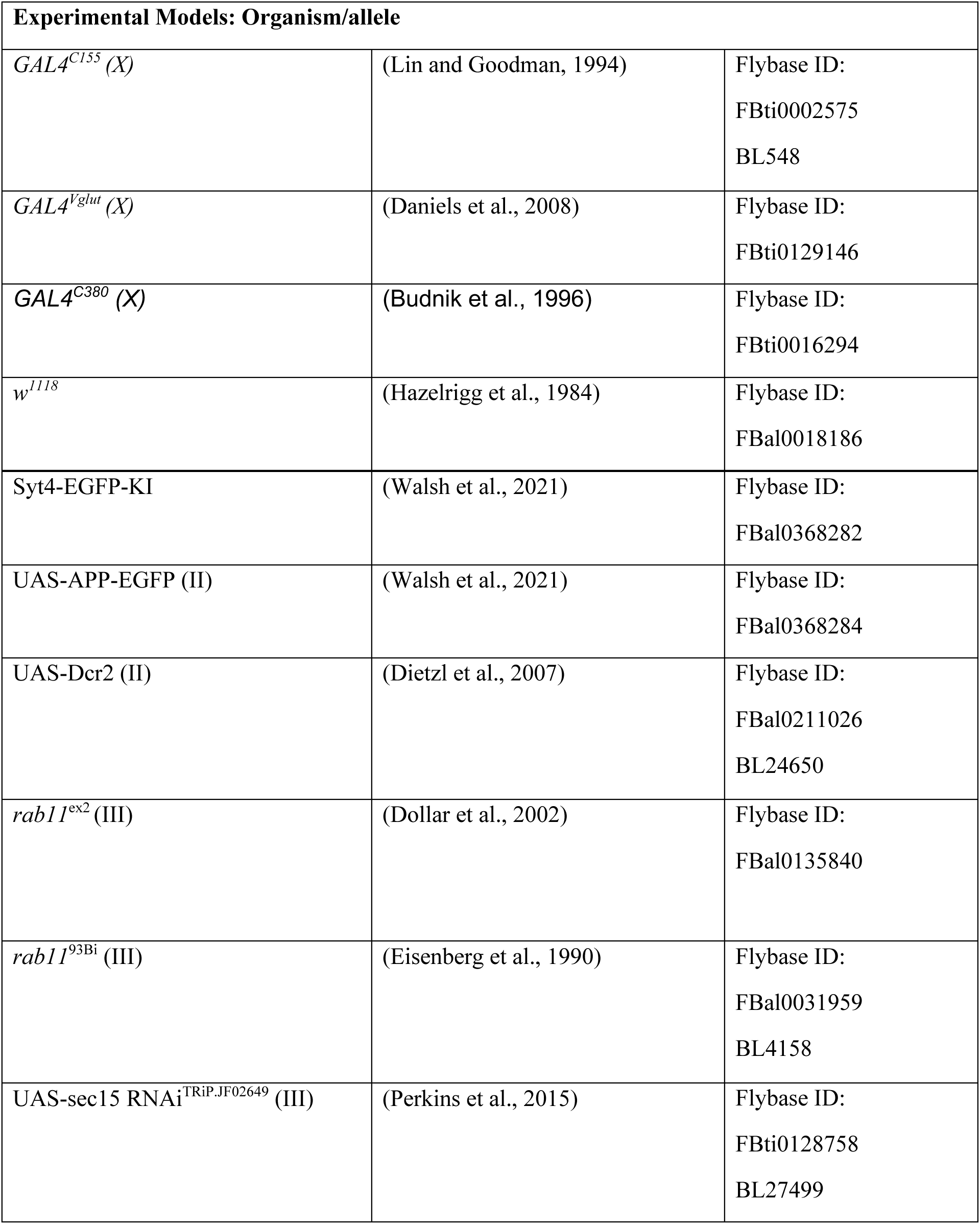
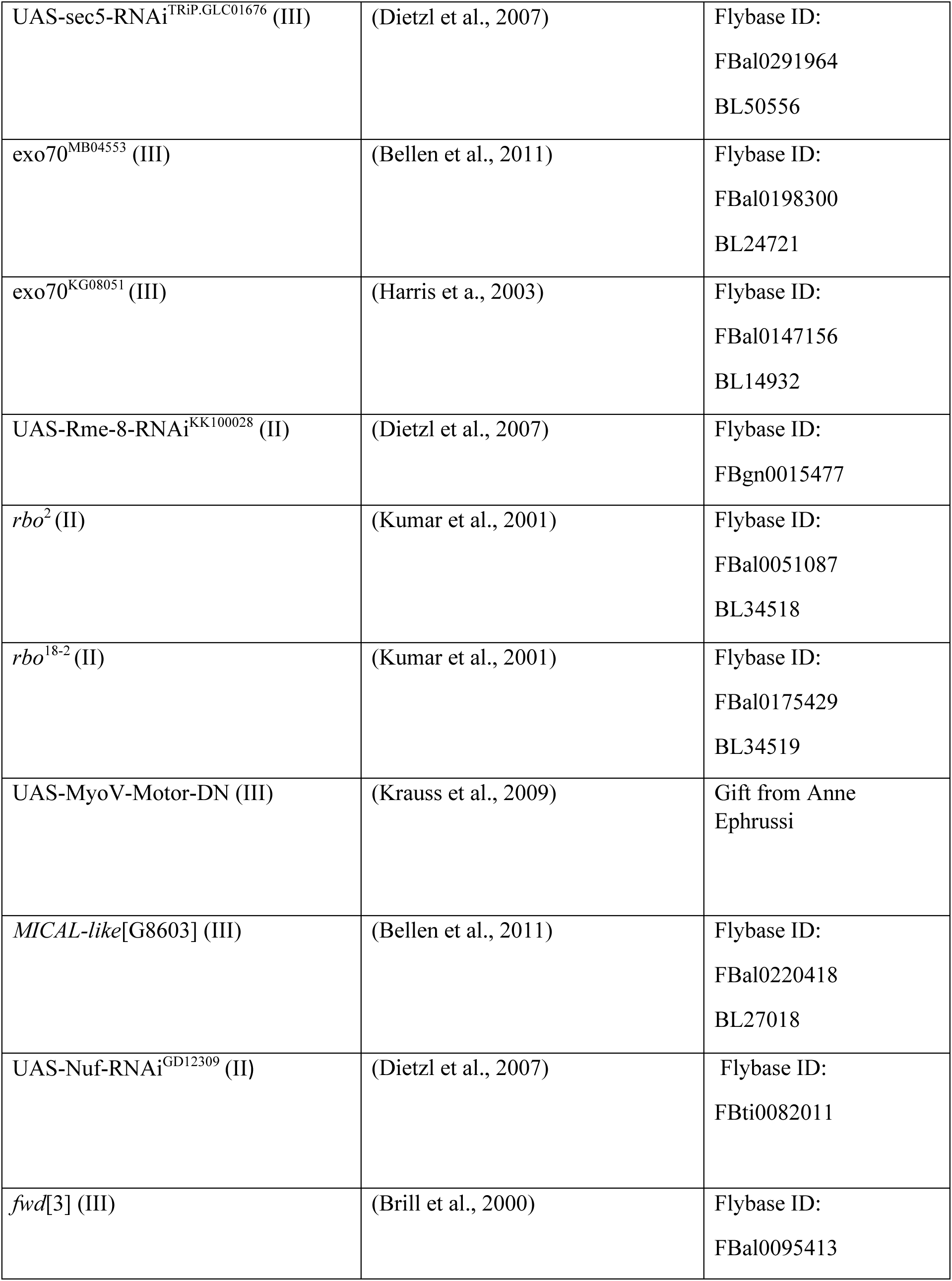
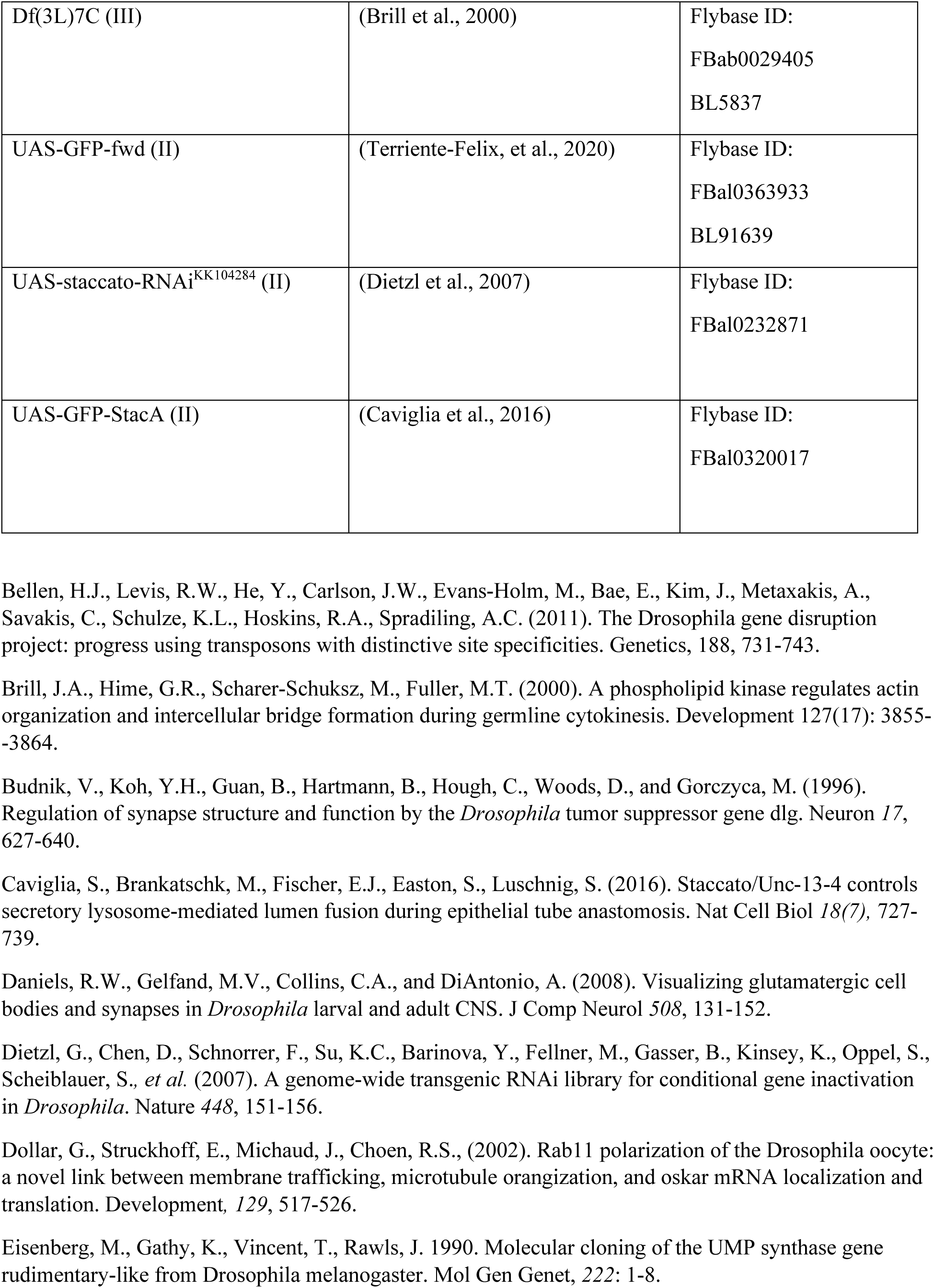
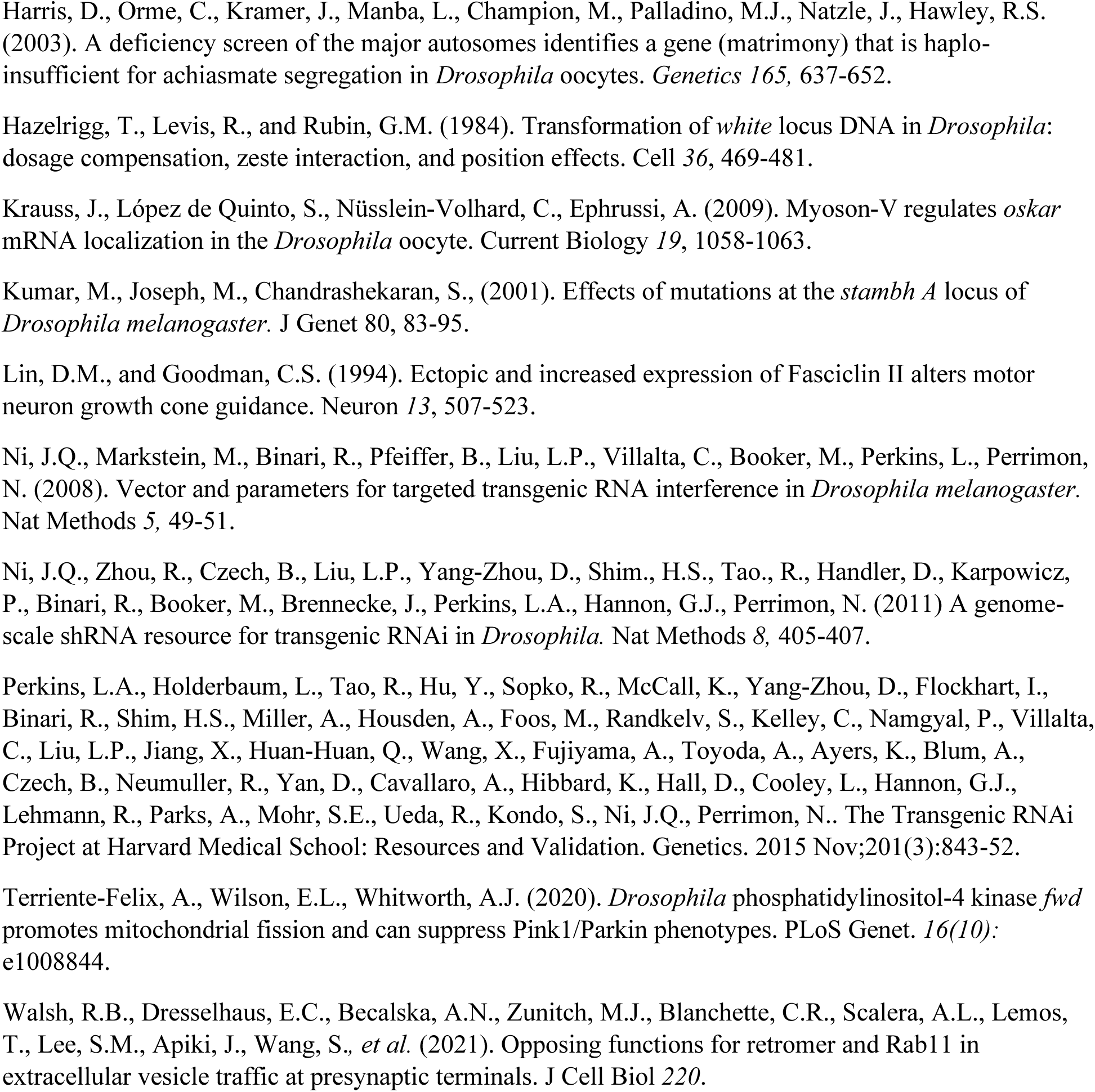
*Drosophila* strains. Bloomington Drosophila Stock Center stock numbers denoted with BL; chromosome for transgene
insertion indicated in roman numerals.

**Table S3.** Antibodies and reagents.

| REAGENT | SOURCE | IDENTIFIER | Concentration |
| --- | --- | --- | --- |
| **Antibodies** |  |  |  |
| α-HRP-488, -RRX, -647 | Jackson ImmunoResearch | | 1:250-1:500 |
| Donkey α-Goat Alexa594 | Invitrogen | A11058 | 1:500 |
| Goat α-Chicken STAR RED | Abberior | 11005 | 1:200 |
| α-Nrg | (Hortsch et al., 1990), DSHB | BP104 | 1:100 |
| α-Nuf | (Rothwell et al., 1998) | | 1:1000 |
| $\alpha$ -Golgin-245 | (Riedel et al., 2016), DSHB | Golgin245 | 1:1000 |

Hortsch, M., Bieber, A.J., Patel, N.H., and Goodman, C.S. (1990). Differential splicing generates a nervous system-specific form of *Drosophila* neuroglian. *Neuron* 4, 697-709.
Riedel, F., Gillingham, A.K., Rosa-Ferreira, C., Galindo, A., and Munro, S. (2016). An antibody toolkit for the study of membrane traffic in *Drosophila melanogaster*. *Biology Open* 5(7), 987-992.
Rothwell, W.F., Fogarty, P., Field, C.M., and Sullivan, W. (1998). Nuclear-fallout, a *Drosophila* protein that cycles from the cytoplasm to the centrosomes, regulates cortical microfilament organization. *Development* 125(7), 1295-303.

